# GARN3: A coarse-grained helix centered technique for RNA 3D structures prediction

**DOI:** 10.1101/2025.07.05.663322

**Authors:** Jhonatan Silva, Johanne Cohen, Daniel Cordeiro

## Abstract

The study of the prediction of three-dimensional structures of RNA (ribonucleic acids) has been increasing over the last decades, especially with advances in artificial intelligence. However, there are still many gaps. Among the known techniques, GARN (Game Algorithms for RNa 3D sampling) has demonstrated good results in molecules with long structures containing hundreds of millions of nucleotides. Nevertheless, the GARN technique also left room for improvements when considering its final 3D structure for predicted molecules, which can be more refined by presenting more elements, also known as pseudoatoms. In this study, we improved the last version of the technique GARN, which is GARN2, enhancing the visualization of the 3D model by adding more pseudoatoms to the helices in our technique GARN3. In GARN3, we also added a machine learning prediction model to improve the scoring calculation, aiming to improve the prediction of new molecules. In our tests, we demonstrated good results when comparing GARN3 with other techniques in the literature. GARN3, in the same way as GARN2, presents better results for large molecules. In our tests, the simulations with GARN3 demonstrated good results, where GARN3 predicted the majority of molecules (approximately 80% of the test set) better than the older versions of GARN.

## Introduction

Ribonucleic acids (RNA) are molecules present in living beings, responsible for functions such as protein synthesis through the translation of the genetic code of individuals, catalyzing biochemical reactions, and performing regulatory roles. Studies reinforce the importance of 3D structures of RNA, for example, in the production of certain drugs, and can also provide the discovery of disease-causing mutations [1].

To obtain such structures with the least probability of errors, experimental techniques are used in the laboratory, which, however, are time-consuming and expensive [2]. For this reason, computational techniques have been developed in recent decades in an attempt to predict such structures. Recently, there are very well-known techniques such as AlphaFold3 [3], which uses artificial intelligence to perform simulations. The technique is currently open source and is available to researchers. However, there are limitations, such as execution time and/or length of the molecule sequence to be predicted.

There are many prediction techniques that benefit from known RNA structures that were obtained by experimental techniques. These techniques are usually developed by taking advantage of physics interactions [4, 5], statistical potentials such as knowledge-based (KB) functions [6–11], or even databases of other molecules to simulate structures of new molecules [12–16]. One of these studies, in which the technique uses statistical potentials, GARN (*Game Algorithms for RNa 3D sampling*) [10, 17], has demonstrated the ability to overcome the time and length limitations of previous techniques, using game theory concepts in its molecular simulations. Game theory is an area of mathematics whose objective is to analyze interactions between two or more elements within a set, where the actions of one element must be taken by always checking the possible actions of the others [18].

In GARN, simulations use a coarse-grained representation based on a 3D graph. It starts with the algorithm input, which is the primary and secondary structure, where the initial 3D graph is generated with the nodes representing the nucleotides of the molecule. Each node of the graph represents one or more nucleotides, relying on the Secondary Structure Element (SSE), such as helices, loops, and *n-way* junctions. The nodes of the graph are linked using a predefined distance according to the SSE type of the node. To generate the position of each node, game theory is applied following this input: players are the nodes of the graph, and strategies are a set of predefined angles in which one player can be linked to another, aggregated by SSE type. Each game aims to generate a possible molecule by moving each player using the given strategy. At each turn, the probability of each strategy to be chosen for each player is updated using two regret minimization algorithms based on machine learning methods. The goal of these regret minimization algorithms is to maximize the gain or minimize the loss in the game by using the payoff of each player.

The payoff for each player in a game’s turn is defined by a KB function, using a well-known intermolecular potential named Lennard-Jones [19]. The inputs of this KB function were defined after several analyses of native molecules as well as the pre-defined strategies of the game. The simulation is finished when several games are generated, using the concept of multi-armed bandit problem [20], where many “arms” (instances) are pulled to play the games, and the game that has the best result is chosen to build the final 3D structure.

Taking into account previous studies for the technique GARN [10, 17], the main difference is that GARN2 is able to predict molecules with 4-way junctions or higher order junctions, whereas GARN can only predict molecules with topology up to 3-way junctions. However, previous GARN studies left some opportunities for improvement related to the prediction of small molecules (usually fewer than 100 nucleotides), where the 3D model presents few pseudoatoms compared to the number of nucleotides in the structure. A possible solution mentioned by the authors is to change the game settings to obtain more refined structures containing more pseudoatoms, that is, visually closer to the structures obtained by experimental techniques.

To overcome the limitations of small-molecule predictions and improve the 3D model of all-molecule predictions, we introduce GARN3, a knowledge-based technique to predict coarse-grained 3D structures of RNA, using statistical potentials with the support of a machine learning model. The main improvement between this study and the previous studies of GARN is the graph that was updated, bringing more elements (pseudoatoms) to the final 3D structure. In previous studies, the SSE helix node aggregated several nodes depending on the helix length, with a maximum of five base pairs per node. With the updated graph, each base pair is represented by one node, regardless of the length of the helix. To present the new graph with more pseudoatoms, the distance between the nodes, the strategies used in the game, and the scoring function parameters were refined and updated.

Furthermore, the scoring function from the previous study only considered static values in the parameters. This study uses a machine learning prediction model to obtain the distances by SSE type from the scoring function. With the new scoring function, we have improved the predictions compared to GARN2, since the machine learning model used in GARN3 considers more characteristics of the molecule than GARN2. In addition, the scoring function can be easily updated in subsequent studies, helping to improve the GARN model even more.

### Related works

To validate the results of the GARN3 technique, we compared its predictions with those generated by other established methods. To this end, we reviewed previous studies that describe RNA 3D structure prediction techniques utilizing primary and/or secondary structure as input, focusing specifically on methods that are available online. For this purpose, there are recent review studies that present a mapping of techniques to predict 3D structures of RNA [21–23].

The techniques found in the reviews are commonly classified according to the type of simulation applied in the technique. The techniques FARNA [4]/ FARFAR/ FARFAR2 [12], MC-Sym [24], vFold [25]/ vFoldLA [15], RNAComposer [13], 3dRNA [26]/ 3dRNA v2 [16] and FebRNA [27] found in the review by Clément et al. [21] are classified as *template-based* (or *fragment-assembly*), which means that RNA homology structures (obtained in experimental techniques) are used to derive others, either entirely or partially. The studies for ModeRNA [28] and RNABuilder [29] techniques are also in this category and are mentioned in the reviews; however, unlike the other studies in this category, ModeRNA requires a 3D structure as input and RNABuilder requires base pair interactions as input.

Another category is *template-free* (*ab initio*), which aggregates techniques that do not use RNA homology structures in the predictions. These techniques generally make use of physics interactions or knowledge-based statistical potentials [30], based on observations after previous analysis of RNA homology structures. In the review by Clément et al. [21], the techniques iFoldRNA [5], NAST [6], Ernwin [8], IsRNA [31]/ IsRNA1 [11], SimRNA [9], RNAJAG [7], RNAJP [7] are classified as template-free. Other techniques found in the review [21] are OxRNA [32], HIRE-RNA [33] and, in the review by Mukherjee et al. [22], the RNA-BRiQ [34] technique is found; however, unlike the other techniques in this category, the RNA-BRiQ and OxRNA techniques require complex inputs not found as primary and secondary structures, HIRE-RNA technique is not available online and BARNACLE [35] is referred in the review [21] as not runnable (with the code avaiable online by the authors).

The last category mentioned in the reviews [21–23] is referred to as *deep learning*, which includes techniques that use artificial intelligence models that incorporate methods such as multiple sequence alignment (comparison between primary and secondary structures to find similarities and extract functions or geometric 2D/3D patterns) and geometric features (angles and distances between atoms) [23]. In the reviews [21–23], the techniques found are AlphaFold3 [3], RhoFold+, trRosetttaRNA [36], DRfold [37], RoseTTAFoldNA [38], PAMNet [39] and epRNA [40]. Another technique found in the reviews [21, 22] is NuFold [41], which is not available online.

## Materials and Methods

Game theory is an area of study on interactions in a set of elements, named players, where the actions of each player influence the other players’ outcomes; the possible actions for players in a game are called strategies [18]. Each strategy chosen by each player in a game applies a reward or a penalty to all the players. Therefore, a strategy may benefit one player more than others. In order to have an equilibrium in the rewards and penalties of all players in a game, a possible approach is called regret minimization. Regret minimization is a widely used decision-making approach for unknown environments, in which we can project many possibilities of a game by applying a scoring function to players after choosing a possible strategy and, in the end, select strategies that provide a good balance of rewards and penalties for all players [18].

In GARN3, the players are pseudoatoms of a molecule, the strategies are the directions where the next pseudoatom can be placed in the 3D space, and the rewards (or penalties) for each player are given by a KB scoring function applied in regret minimization algorithms. In the context of this study, the KB function reflects the best distance between 2 pseudoatoms, considering characteristics such as SSE type and the length of molecules. Unlike GARN/GARN2 [10, 17] techniques, for the KB function, in GARN3 the best distance is retrieved by a machine learning model instead of static values in the algorithm, providing better results by considering molecule characteristics such as SSE types present in the molecule, length of the molecule, length of the helices in the molecule, among other characteristics that are listed in this section. The input for GARN simulations is the primary and secondary structure of a molecule, and the output is a file with the 3D coordinates for the same molecule.

In order to enhance the GARN model by reducing the granularity, we propose a new refined 3D model containing more pseudoatoms. GARN configurations were updated using information derived from a dataset of 580 molecules. This dataset provides information on the primary, secondary, and 3D structures of these molecules. A detailed statistical analysis of the frequency distribution of the angles and distances between atoms in the molecules is presented and will be used to motivate a proposal for a new set of strategies for the game and new parameters for the scoring function that are better suited for the proposed representation model with less granularity.

The algorithms for regret minimization applied in the GARN3 technique are presented in this section. At the end of each molecule simulation, after rounds of a game, a sorting criterion needs to be applied to identify which structure is the best to be used as the final one. Therefore, the sorting criteria for the generated structures are listed in this section. Also, taking into account the players and the 3D representation, this section describes the relation between players and an SSE and how the final 3D model is represented.

### Molecules dataset

In order to obtain a knowledge-based configuration for the games, a well-known protein database named RNA FRABASE [42] was used to collect primary and secondary structures. For 3D structures, the RCSB Protein Data Bank [43] was used. This set of molecules was used to calculate the distance and angle between the elements of the structures and the length of each player to determine the parameters for the scoring function, define strategies, and also evaluate the predictions.

The sample consists of 580 molecules for data analysis and 22 molecules as a test set to use as a comparison for the predictions in this paper. This amount was obtained considering different aspects:

- Several molecules with almost identical SSE and 3D models were not considered because they could be negatively influenced if repeated multiple times. The similarity between these molecules was almost identical and was measured using the RMSD calculation [44].
- Molecules with an incomplete 3D model were not considered because the distances and angles between the atoms in the nucleotides led to an inconclusive result.

For the GARN model, we considered important to understand the behavior when predicting small (less than 100 nucleotides), medium (between 100 and 1000 nucleotides) or large molecules (more than 1000 nucleotides) of some specific SSE types, which are 2-way, 3-way, n-way (having *n >* 3) and molecules with pseudoknot. The list of molecules used for the analysis and also to generate the machine learning model is presented in S1 Appendix, and the test set is presented in S1 Table.

### Representation of the molecules

Except for the helices, the other elements in the final three-dimensional representation of the molecule follow the representation present in GARN [10].

- Each helix is represented by *n* players, where *n* is the number of base pairs in this helix. As mentioned earlier, this differs from the GARN/GARN2 model, which has a variable number of players relying on the junction’s length. An example is shown in Fig 1, where the molecule has more beads in the GARN3 model than in the GARN/GARN2 model.
- Terminal loops and 2-way junctions of any type are represented by 1 player.
- 3-way junctions are represented by two nodes: 1 to support the base from where the previous player is coming from, and the other to support the player in the next two 1-way junctions (1 for each branch of the 3-way junction).
- N-way junctions having *n* higher than 3 follow are represented by the number of players following the equation:

**Fig 1.**
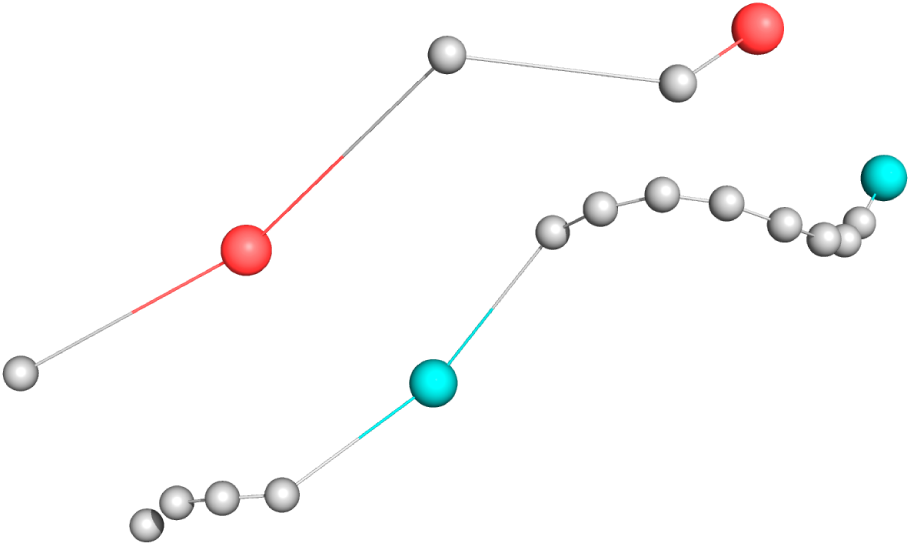
Representation of a GARN/GARN2 3D molecule model, in orange, compared to the GARN3 3D model, in cyan. This model is generated using the native PDB file from the RCSB database. The PDB ID for this molecule is 1MNX.

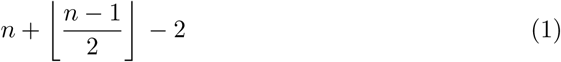

Each of these players is represented as a node in a three-dimensional graph. A two-dimensional graph of a given molecule is illustrated in Fig 2.

**Fig 2.**
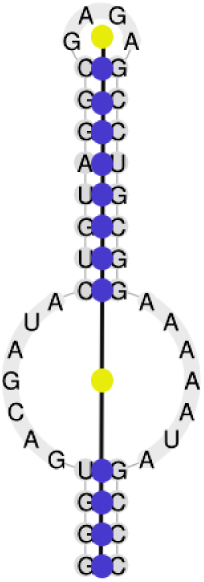
Representation of a GARN graph of a predicted molecule in 2D. Each base pair is represented by one node, while the 2-way junction is represented by one node, as it is in the study of GARN2 [10]. The PDB ID for this molecule is 1MNX.

Another aspect related to the 3D representation is the length of the edges, which is the distance between the player elements. For nodes in helices that represent a base pair, the edges have a length of 2.4 Å, where this value is present in previous studies [10, 45]. For nodes from other SSE types, this study uses the same lengths as present in the GARN [10] study.

### Players and Strategies

At each turn of the game, for each player of the molecule, a strategy is chosen from a set of possible strategies. The strategies for the GARN model are directions in the 3D space where the player (pseudoatom) can move towards the next players. Similarly to GARN/GARN2, the order of players in GARN3 is defined by applying a depth-first search starting from the larger junction, which can be seen in S1 Fig). These strategies are chosen until the last turn is finished, then the best turn will be chosen to represent the prediction.

Also, similarly to GARN/GARN2 [10, 17], the set of possible strategies is available in an interval of 30° or 60°. However, to avoid having too many strategies unnecessarily, the set of strategies for each SSE type is classified by junction types. That way:

- For 2-way junctions, there are 2 situations. In both cases, the angle interval is 30°.

**–** For bulges (internal loops in which one of the unpaired sides has length zero), the strategies range from 30° to 120°.
**–** In any other case, strategies range from 30° to 60°.
- For 3-way or high-order junctions, strategies vary from 0° to 180° by an interval of 60°.

In order to update the strategies for GARN3, we analyzed angles from structures within our molecules database. To perform these analyses, the molecules of a native all-atom representation obtained from the Protein Data Bank (PDB) were transformed into the GARN3 model. After that, calculations on the angles between two players were made considering different aggregations of SSE types. The metric used to measure the angles in the histograms is radians. In Fig 3, the frequency of angles is represented in two separate contexts: on the right side, only molecules with 3-way junctions or higher-order junctions are allowed, and on the left side, any other molecule with helix, terminal loops, and 2-way junctions is allowed. After the analysis of angles in the histograms, the strategies for the players of helix SSE type were updated as follows:

- For molecules which contain only Terminal (stem-loop), Helix, and 2-way junctions: 0°– 30°
- For molecules with 3-way or higher-order junctions: 0°– 60°

**Fig 3.**
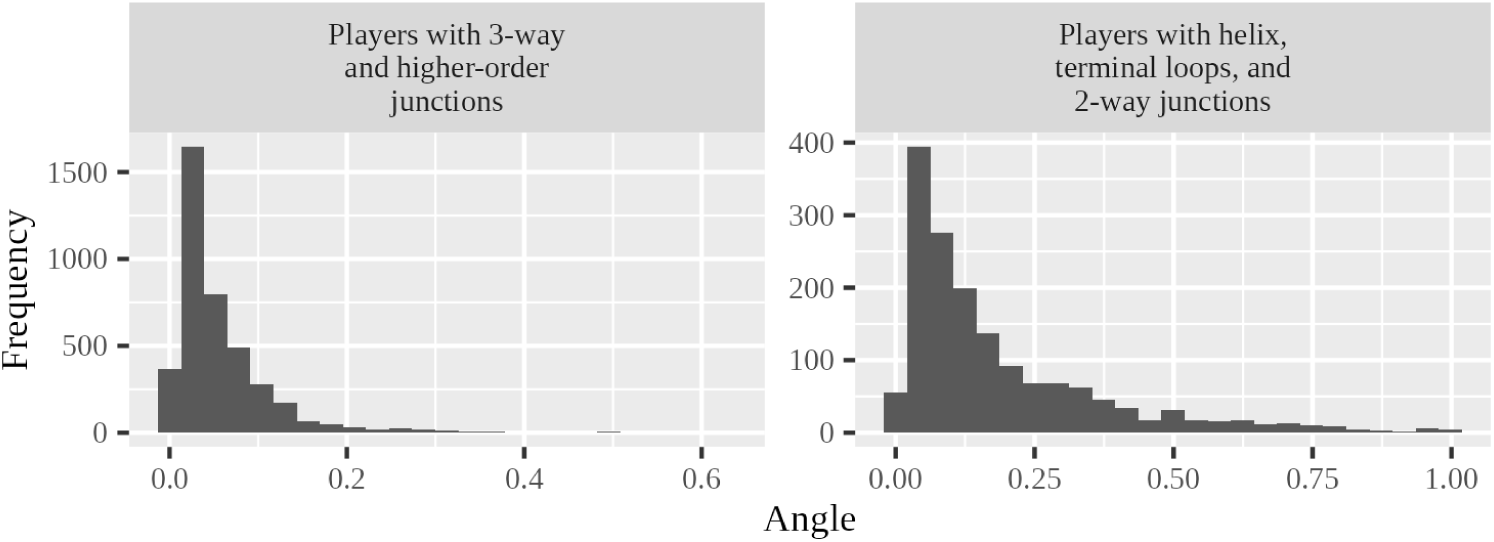
Strategies for players of helix SSE type.

### Algorithms for Regret Minimization

The regret minimization algorithms EXP3 [46] and UCB [47] were implemented in GARN3. Considering the analysis in the study of GARN [17], the application of EXP3 resulted in better predictions for molecules with 3-way or higher order junctions and UCB otherwise. In this paper, it is analyzed whether the prediction comparisons classified by molecule type using a specific regression minimization algorithm follow the same pattern as the GARN [10] study; in other words, if EXP3 is still better for 3-way or higher-order junctions and UCB for the other cases.

Using the same approach as from GARN/GARN2 [10, 17], the implementation of GARN3 regret algorithms in game simulations is as follows:

1. At the beginning of each game, all the players choose a possible strategy and compute a molecule build using those strategies;
2. After each turn of the game, scores and probabilities are updated for each of the players;
3. If, at any moment of a game, the molecule is invalid, the game is canceled, and the players are penalized. In GARN3, a molecule is considered invalid when two edges intersect.

### Scoring

To define the probability of each possible strategy, after each turn of a game, a scoring function is applied, considering a KB approach (knowledge-based) on the distance between atoms in a molecule. In this KB function, a probability is calculated using a function based on the Lennard-Jones function [19], defined as

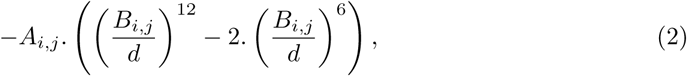

where *A* and *B* are defined according to the type of player (helix or junction).

The parameter *B* is used in two different ways: a static value that depends on the SSE type, defined by the analysis of the molecules in the database; and a dynamic value that depends on many parameters, using a machine learning prediction model.

### Scoring Static Parameters

In order to update the scoring parameters, we performed an analysis on distances between atoms in the molecules from the database in this study. To perform these analyses, the molecules of a native all-atom representation obtained from the PDB files were transformed into the GARN3 model, similar to the analysis to update the strategies. Afterwards, the distance calculations were done considering different aggregations on the SSE types. For each updated score, the value represents the best distance considering a histogram; in other words, the value with the highest frequency in the histogram. The metric used to represent the distances between atoms or structures in this paper is angstrom (Å), where 1 Å = 10*^−^*^10^*m*.

For an interaction between two players of helix SSE type, the results are represented in Fig 4. On the left side of the image, only distances from players in molecules with 15 or fewer players (considering the total number of players in the molecule) are used. On the right side of the image, any other molecule with more than 15 players is used.

**Fig 4.**
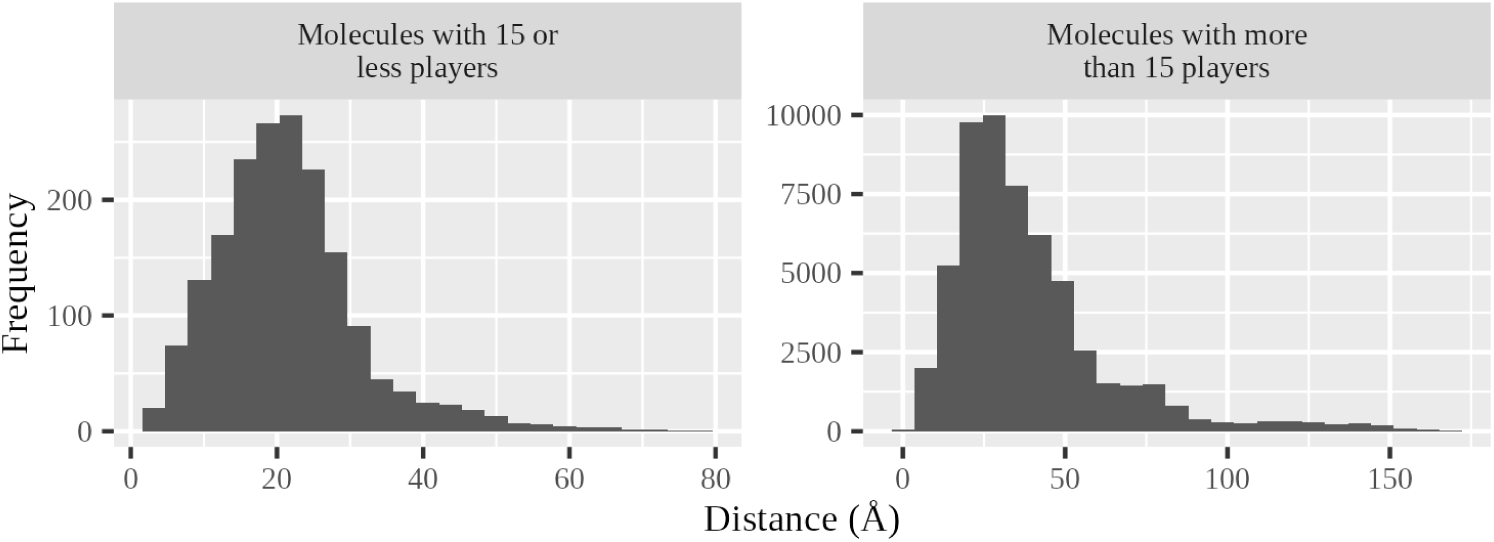
Scores for interactions between two helix players.

For an interaction between players of the helix and 2-way junction SEE type, the results are represented in Fig 5. On the left side of the image, only molecules with a single stream; on the right side, molecules with 2-way junctions.

**Fig 5.**
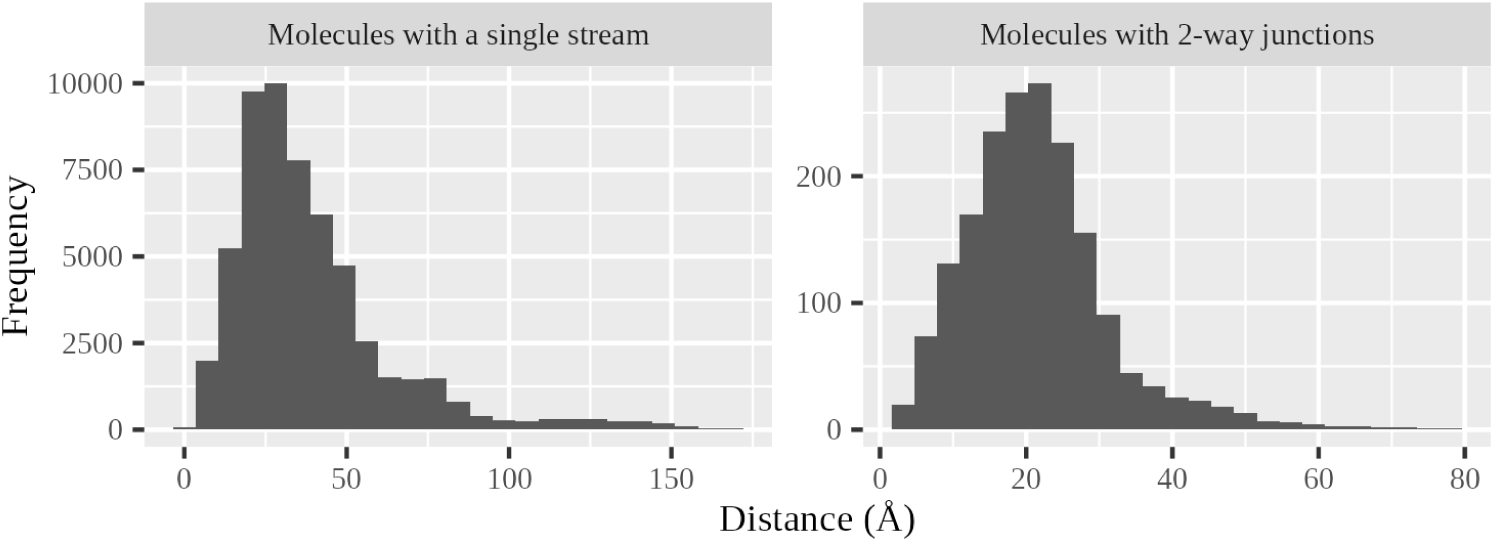
Scores for interactions between a helix player and an unpaired stream player.

After the data analysis on molecules was completed, some of the scoring function parameters were updated. These are related to the Helix SSE type that interacts with other SSE types. The updated static parameters are as follows:

- Two Helix players interacting: 18 Å, if the molecule has at most 15 players; 23 Å otherwise.
- A Helix and an unpaired stream player: 14 Å if the unpaired is a single strand; 26 Å if it is a terminal loop.
- A Helix and a 2-way junction player: if the molecule has only Terminal, Helix, and 2-Way SSE types, 11 Å; or 22 Å otherwise.

For the other interactions between SSE types, this paper follows the previous settings of the GARN2 study [10].

### Machine Learning Prediction Model

In order to obtain more accurate distances to be used in the GARN3 scoring function, different algorithms were tested for a machine learning regression model. They are: Gradient Boosting, Decision Tree, Multilayer Neural Networks, LASSO, Ridge, Random Forest, Linear Regression, k-nearest neighbors, and SVM. The features used in these algorithms are described below. They represent characteristics of the SSE of a molecule, where some of them consider the entire molecule and others consider only the interaction between two elements; these elements are players of a GARN model.

To train and test these models, the molecules in the set were first transformed into GARN players, so they could be used as input files. Therefore, the database consists of a set of GARN games, where the coordinates of these players correspond to the coordinates of the molecules obtained from the RCSB Protein Data Bank. The cross-validation method *k-fold* was applied using 10 folds, which is a popular choice and is likely to be accurate [48].

To define the features, research was done on the literature in order to understand the possible reusability of aspects of similar studies. Some recent studies reviewing the state of the art in RNA prediction with machine learning [49–52] explain the advances; however, the results of these models are not similar to the results needed in this study. In addition, the input of these models considers different aspects of the context of this study.

An investigation of characteristics that could be relevant to determining the distance between two GARN players was carried out, considering the SSE types and the placement of the players in the GARN3 model, which is represented as a graph. The interactions in this machine learning model are always between 2 players: the current player, which is the player whose strategy is being chosen at the moment, and the next player, which can be any other player in the game depending on the iteration; since GARN3 algorithm applies a scoring function to all players in each game turn, the next player will be each player of the molecule except for the current player. Afterwards, the relevant features defined are as follows:

1. SSE type of the current player in the game;
2. SSE type of the next player in the game;
3. Minimum distance between these 2 players, considering that GARN3 model is represented as a graph. For that, the Dijkstra algorithm [53] was used.
4. Quantity of base pairs inside the n-way junction, in case there is an interaction between 2 base pairs at the n-way junction. Otherwise, this value is zero;
5. If players are placed at the same n-way junction, this feature has a value of 1; otherwise, its value is zero;
6. If current and next players are pseudoknots interacting (kissing loops), this value is 1; otherwise, its value is zero;
7. If the current player and next player are a helix and a bulge, this value is 1, and 0 otherwise;
8. Largest junction quantity in the molecule. For example, if the molecule has only base pairs and a terminal-loop, this number is 1. If the molecule has a 2-way junction, this number is 2, and it follows the same logic for higher-order n-way junctions;
9. If this is an interaction between 2 players that are n-way junctions, having *n >* 1, captures the length of the smallest side of this junction; otherwise, this value is −1;
10. If this is an interaction between 2 players that are n-way junctions, having *n >* 1, captures the length of the largest side of this junction; otherwise, this value is −1;
11. Total of players considering the entire molecule.

The output of this regression model is the best distance between two players, given the features as input.

### Sorting criteria

After the rounds of a game, the structures are generated, but at this point, it is not yet defined which structure is the best to be chosen by the GARN3 algorithm. In the GARN2 study [10], 3 approaches were presented: Sorting by global score, where the sum of all players’ scores is taken; Sorting by minimum score, where the molecule with minimum score would be the best one considering that it has less penalization, or the penalization was not relevant; and sorting by maximum distance, where the maximum distance between the players is taken.

From these 3 criteria, the maximum distance is applied in this work, considering that it was validated as the most relevant in the GARN2 study. This sorting criterion will also be validated in this study during the simulations from the test set.

### Evaluation of molecules

For each of the molecules, a widely used metric in the literature named RMSD (Root Mean Squared Deviation) [44] was used. This metric indicates the mean distance between the two molecules, considering each of the elements inside. The RMSD is defined by

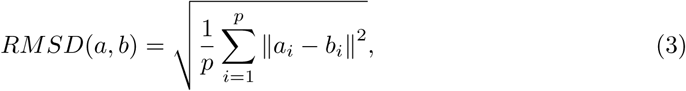

where *a* and *b* are two molecules, *p* is the number of elements (which is a number of players in GARN) *i* in the molecule and *a_i_* and *b_i_* is the position of the player *i* in the molecules *a* and *b*.

In the previous section, several techniques found in the literature are mentioned, which aim to predict 3D structures for RNA molecules. In order to improve the GARN3 simulation results and further discussion, we selected 12 techniques from the literature, having 4 template-free techniques (iFoldRNA [5], NAST [6], SimRNA [9] and IsRNA1 [11]), 6 template-based techniques (FARFAR2 [12], RNAComposer [13], VFoldLA [25], 3dRNA [16], FebRNA [27] and MC-Sym [24]) and 2 deep learning techniques (AlphaFold3 [3] and trRosettaRNA [36]). Although GARN and GARN2 generate molecules with fewer beads than GARN3, GARN2 is used for comparison once it is closely related to this study.

## Machine Learning Model Results

Some metrics were used to define the machine learning prediction model. In order to define the features, an important method to assist in the definition process is the SHAP (SHapley Additive exPlanations) [54]. This method describes numerically the importance of each feature used as input in the prediction model. After the refinement of features with the support of SHAP method, the most relevant ones are described in the following section.

After defining the features of the model, some analysis was performed to identify an appropriate machine learning algorithm to obtain the distances for the scoring function. Many metrics are available in the literature, but MAE and RMSE are among the most relevant to assess model performance [55, 56]. The MAE (Mean Absolute Error) calculates the average error difference between predicted and actual values, regardless of outliers or high values; Therefore, if the MAE is 10 in our model, it means that we have an average of 10Å of difference between the predicted and actual values. The RMSE (Root Mean Squared Error) metric is the root of MSE (Mean Squared Error), and MSE calculates the squared error, providing more weight to large errors. The difference in using RMSE instead of MSE is the scale; MSE is squared, whereas RMSE returns the result of MSE to the original label scale.

In our tests, Gradient Boosting demonstrated better performance over the other algorithms, as illustrated in Fig 6. After running the k-fold cross validation, we retrieved the values for the metrics MAE as ≈ 8Å, the RMSE as ≈ 11Å, and the *R*^2^ (R Squared) metric was also considered. *R*^2^ is a real value between 0 and 1 that represents the difference between the predicted data and the real value. The closer to 1, the better the model. For the model in this study, *R*^2^ has the value ≈ 0.7.

**Fig 6.**
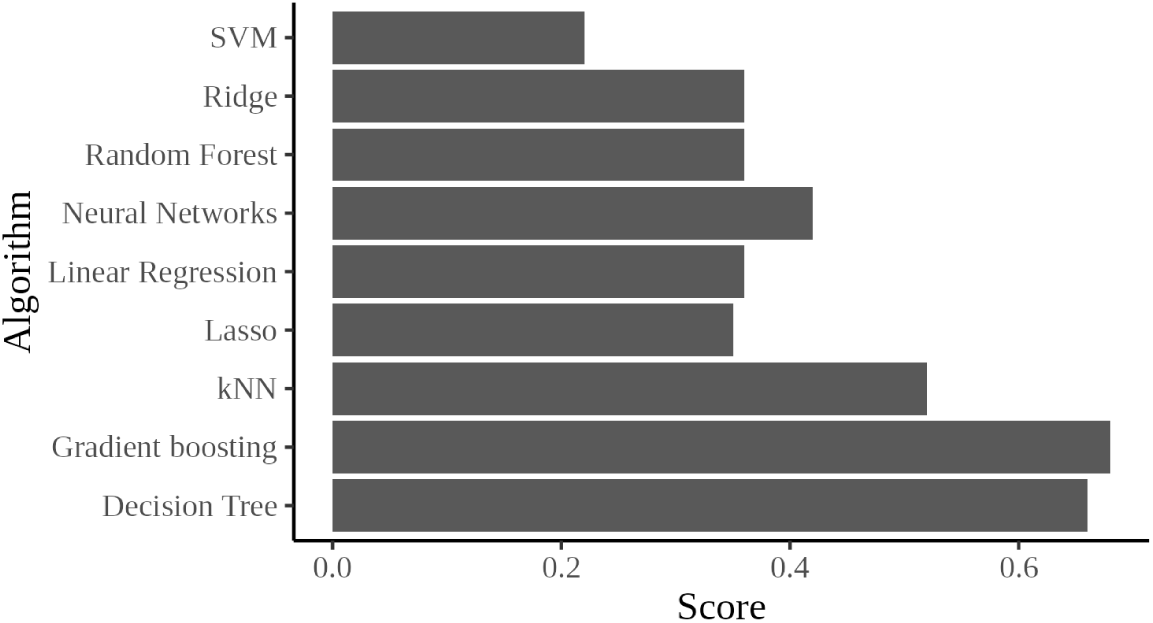
Comparison of common machine learning regression algorithms using a dataset suitable for predicting distance between GARN players. The score described is R^2^.

An analysis of the error on prediction was also performed about the error on the prediction by the Gradient Boosting algorithms using our database as input, as illustrated in Figure 7. In this figure, it is possible to see the predicted distances compared to the actual (expected) distances, split by a line in the middle, where a correlation is visible once there is no significant difference between these predicted and actual values. This comparison is represented by the *R*^2^ value.

**Fig 7.**
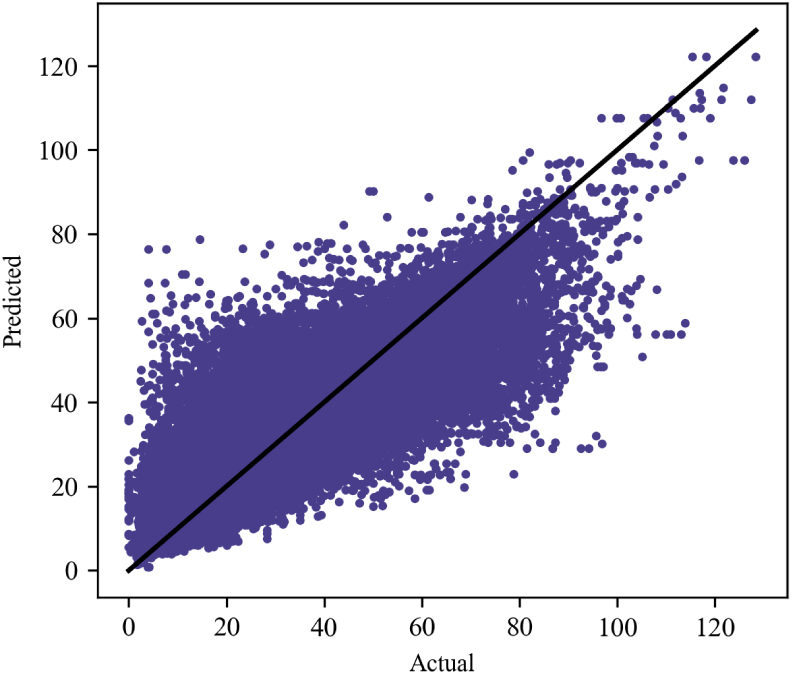
Comparison of real and predicted values using the Gradient Boosting regression model.

Since the use of machine learning in the scoring function differs from previous GARN studies [10, 17], it is important to understand whether the new approach works properly. Hence, simulations were performed using the database of molecules in this study applying the scoring function using static parameters and parameters retrieved from the regression model. The simulation results are aggregated in Fig 8, for all molecules in the test set. The usage of static parameters did not return results for some of the molecules, and they are not present in the figure. Even if the use of dynamic parameters from machine learning is not entirely better than the use of static parameters, it is better for most of the molecules predicted if RMSD is taken into consideration. In addition, there are some situations where the molecule could not be predicted with static parameters, but was possible with parameters by the machine learning model, as seen in Fig 8. Therefore, the simulations in the next section use parameters from the machine learning model.

**Fig 8.**
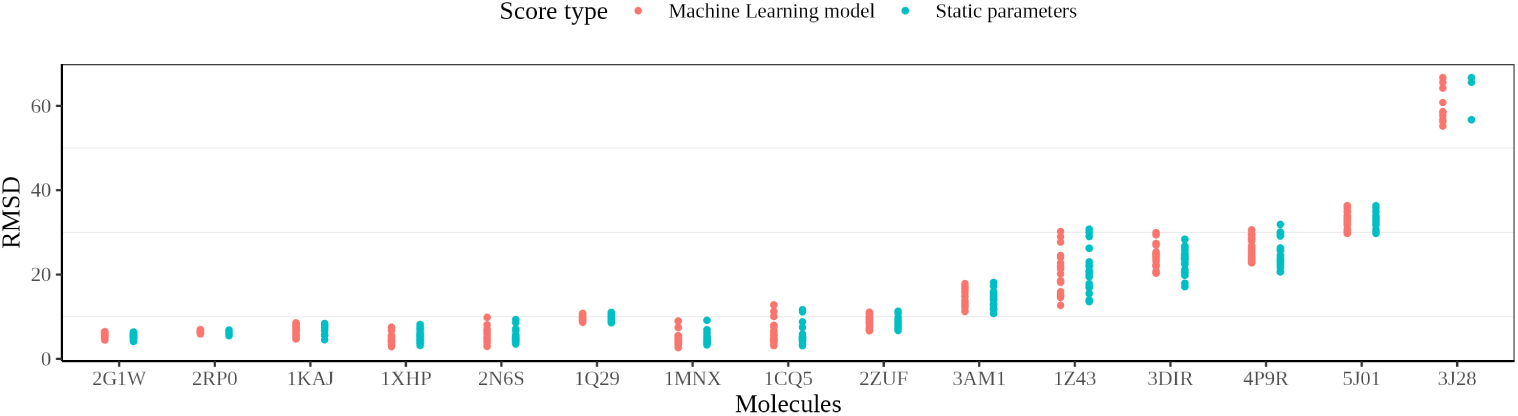
Comparison between simulations using scoring function with static parameters and scoring function with machine learning model. The molecules not present in this graphic were not able to be obtained using the static parameters.

## Simulation results

In this section are presented the results of the experiments, reflecting the best results of the molecules simulations (i.e. results with the lowest RMSD), using GARN3 applying the 2 regret minimization algorithms, EXP3 and UCB. Afterwards, the comparison of results between GARN3 and other techniques, as well as the execution time of these molecules, and the results when sorting and classifying the best molecules by the end of the games of a molecule is also described. For the molecules in each of the techniques used for comparison, 20 simulations were performed.

After updating the game configurations with the new score parameters and strategies, the simulations were run using the molecules database in this study. Later, the results are described and compared with other techniques in the literature. Considering the number of techniques compared in this study, the results of the comparison with other techniques are divided into 3 categories for better visualization of tables and figures, having techniques with template-based, template-free, and deep learning modeling. In addition, in most tables and figures in this section, the molecules in the test set are also divided into categories, based on the SSE types, as follows:

- **2-way junction molecules**, whose junctions have a maximum of 2 ways;
- **3-way junction molecules**, whose junctions have a maximum of 3 ways;
- **N-way junction molecules**, which have at least 4-way or high-order junctions;
- **Pseudoknot molecules**, which have at least 1 pseudoknot.

When requesting simulations for the 3dRNA web server, 3dRNA-Lib2 was used for the predictions, the other parameters being set as default. For all the techniques tested, the same secondary structures used in GARN simulations were provided as input, except for those that do not allow. In this scope, the ones that do not allow are trRosettaRNA and AlphaFold3. NAST simulations were run with 5000 as the number of steps parameter, and IsRNA1 with 20,000,000 steps (the default). VFoldLA was run using all default parameters.

For techniques that do not provide a web server, the simulations were run using a local machine with an Intel Core i7-12800H 12th generation 2400 MHz processor and 64 GB of physical RAM memory.

### Regret minimization algorithms

As mentioned before, the regret minimization algorithms applied in GARN3 are UCB and EXP3. The motivation for using UCB was to understand how the behavior is because, in the GARN study [10], smaller molecules are better predicted using UCB. In the GARN3 model, because there are more players (pseudoatoms) in the final 3D structure for the same molecules, UCB could behave slightly differently.

From the molecules tested, smaller ones, usually with a more simple topology with a maximum of 2-way junctions, demonstrated better results when the simulations were run using UCB. Considering the set, 3 of the 5 molecules were better evaluated with UCB. In addition, from the molecules with pseudoknots, even though their difference was smaller (less than 1 Å), the results were also better evaluated with UCB.

Taking into account larger molecules, with 3-way or higher-order junctions, the results were shown to be better when using EXP3. Another point to consider was the time to predict seen in, for example, the molecule 1Q29, which has 3-way junctions: it took several hours, approximately 20h, to predict the results with UCB, while the usage of EXP3 returned better results in less than 1 minute. All results are listed in S4 Table.

### Simulations from other techniques

Unlike all others, the comparison with GARN2 was achieved with a different approach due to the considerably smaller number of pseudoatoms in the final 3D structure. Using the original PDB molecule, a GARN2 model is generated, with fewer pseudoatoms than GARN3. Using the same molecule, a GARN3 model is generated. The GARN2 model is used to compare the GARN2 simulation results, whereas the GARN3 model is used for the GARN3 results, as well as for the other techniques in the literature; For the other techniques, the output model of the technique (usually a PDB file) is used as input in the GARN3 algorithm to generate a GARN3 model. In Fig 9, both models are exemplified, having the original molecule on the left side, the GARN2 model in the upper right corner, and the GARN3 model in the lower right corner. For all other techniques, the GARN3 model was used for comparison. There is an example of comparison in S2 Fig with the molecule 1XHP, where the best structure predicted by each of the techniques is represented, compared to its native structure.

**Fig 9.**
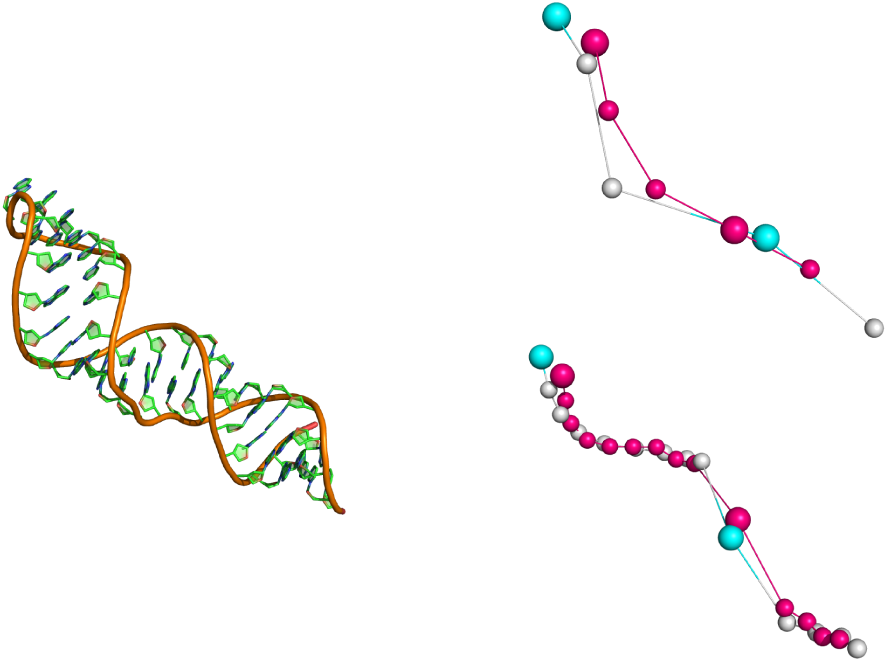
Comparing RMSD with GARN2 and GARN3 models, where GARN2 has less pseudoatoms. Both darker models in the right corner represent the predicted structure, and the lighter models are generated using coordinates from the molecule’s native structure (obtained in laboratory).

After running the simulations of GARN3 and other techniques, the results of minimum and maximum RMSD by molecule and technique are presented in Table 1, considering only template-free modeling techniques. In this table, all molecule simulations present the best result considering EXP3 or UCB as the regret minimization algorithm, and the scoring parameters were obtained from the machine learning model.

**Table 1.** Comparison with template-free modeling techniques. The model for comparison in GARN2 was different from the others due to the number of pseudoatoms in its predictions, which is lower than GARN3. The highlighted values in each row aim to help the visualization of the technique with lower RMSD for the molecule.

| Molecule | Type | Length | RMSD | iFoldRNA | NAST | SimRNA | IsRNA1 | GARN2 | GARN3 |
| --- | --- | --- | --- | --- | --- | --- | --- | --- | --- |
| 1XHP | 2-way | 32 | Min | 1.13 | 8.51 | – | 1.12 | 2.15 | 2.90 |
|  |  |  | Max | 1.13 | 8.51 | – | 1.43 | 4.85 | 7.49 |
| 1MNX | 2-way | 42 | Min | 2.99 | 9.84 | 2.42 | 1.47 | 6.96 | 2.67 |
|  |  |  | Max | 15.4 | 9.84 | 5.35 | 3.39 | 9.56 | 8.96 |
| 1CQ5 | 2-way | 43 | Min | 2.93 | 9.18 | – | 3.4 | 2.62 | 2.46 |
|  |  |  | Max | 10.2 | 9.18 | – | 5.58 | 8.68 | 10.48 |
| 2RP0 | 2-way | 27 | Min | 2.59 | 4.59 | 0.897 | 4.21 | 5.51 | 5.42 |
|  |  |  | Max | 2.59 | 4.59 | 1.58 | 7.47 | 11 | 6.8 |
| 2N6S | 2-way | 36 | Min | – | 6.85 | 2.10 | 2.05 | – | 2.97 |
|  |  |  | Max | – | 6.85 | 3.64 | 2.05 | – | 9.83 |
| 1Q29 | 3-way | 41 | Min | 7.28 | – | 2.00 | 2.05 | 4.20 | 8.67 |
|  |  |  | Max | 8.68 | – | 6.87 | 2.05 | 10.2 | 10.8 |
| 3DIR | 3-way | 174 | Min | 13.7 | 24.1 | – | – | 16.2 | 14.1 |
|  |  |  | Max | 21.0 | 24.1 | – | – | 29.9 | 27.19 |
| 4P8Z | 3-way | 188 | Min | 18.5 | 15.9 | – | 23.3 | – | 19.6 |
|  |  |  | Max | 25.2 | 15.9 | – | 23.3 | – | 29.0 |
| 3AM1 | 3-way | 81 | Min | 11.4 | 8.29 | 8.23 | – | 18.4 | 11.2 |
|  |  |  | Max | 16.5 | 8.29 | 14.4 | – | 19.3 | 17.8 |
| 4RZD | 3-way | 102 | Min | 12.5 | – | – | 1.22 | 8.12 | 11.3 |
|  |  |  | Max | 16.0 | – | – | 14.1 | 15.8 | 18.4 |
| 4QKA | 3-way | 122 | Min | 10.4 | 73.2 | – | 9.93 | 12.6 | 11.6 |
|  |  |  | Max | 12.4 | 73.2 | – | 9.93 | 23.0 | 23.6 |
| 1Z43 | 3-way | 101 | Min | 4.22 | 20.3 | – | 14.8 | 11.0 | 12.7 |
|  |  |  | Max | 12.7 | 20.3 | – | 14.8 | 29.3 | 30.2 |
| 4P9R | 3-way | 189 | Min | 27.3 | 29.9 | – | 14.5 | 18.5 | 17.72 |
|  |  |  | Max | 30.1 | 29.9 | – | 14.5 | 27.7 | 30.6 |
| 4OQU | n-way | 97 | Min | 10.7 | 26.2 | 5.32 | – | 10.5 | 10.5 |
|  |  |  | Max | 14.3 | 26.2 | 5.32 | – | 20.7 | 19.8 |
| 4QK8 | n-way | 124 | Min | 11.2 | 18.7 | 13.2 | – | 15.5 | 11.1 |
|  |  |  | Max | 14.5 | 18.7 | 16.5 | – | 23.4 | 26.6 |
| 5J01 | n-way | 418 | Min | 54.1 | 230.0 | – | – | 35.1 | 29.7 |
|  |  |  | Max | 59.2 | 230.0 | – | – | 53.1 | 36.3 |
| 3J28 | n-way | 1533 | Min | 97.8 | 77.0 | – | – | 58.9 | 55.2 |
|  |  |  | Max | 99.8 | 77.0 | – | – | 78.6 | 66.7 |
| 1C2W | n-way | 2904 | Min | – | – | – | – | 69.1 | 66.68 |
|  |  |  | Max | – | – | – | – | 85.4 | 78.1 |
| 2NBX | n-way | 108 | Min | – | 22.7 | 9.75 | 17.2 | 58.1 | 15.5 |
|  |  |  | Max | – | 22.7 | 16.7 | 17.2 | 74.4 | 24.6 |
| 2G1W | pseudo-knot | 22 | Min | 2.64 | 2.76 | 2.05 | 2.14 | 5.81 | 4.46 |
|  |  |  | Max | 2.64 | 2.76 | 2.90 | 2.14 | 11.4 | 6.44 |
| 1KAJ | pseudo-knot | 32 | Min | 3.26 | 4.25 | 4.12 | 4.26 | 6.74 | 4.71 |
|  |  |  | Max | 3.26 | 4.25 | 4.21 | 4.26 | 10.6 | 8.53 |
| 2ZUF | pseudo-knot | 78 | Min | 11.7 | 10.2 | – | 15.4 | 7.07 | 6.64 |
|  |  |  | Max | 16.2 | 10.2 | – | 15.4 | 12.3 | 11.1 |

For 2-way molecules and also for pseudoknots, where these molecules do not have a large length, SimRNA and iFoldRNA demonstrated good results. Taking into account a comparison between GARN2 and GARN3, GARN3 predicted the molecules better in 4 of the 5 molecules. For 3-way molecules, iFoldRNA demonstrated good results. However, the SimRNA technique was not able to predict most of the molecules in this category, nor even when trying to predict higher-order junctions. For 4-way or higher-order junctions (n-way), where molecules are easily found to have more nucleotides, GARN3 demonstrated better results in most of the molecules.

For simulations on template-based modeling techniques, Table 2 presents the results of simulations for each molecule tested. Due to the number of techniques that cause the table to be excessively large, MC-Sym and VFoldLA techniques are not presented. This does not cause any disruption in verifying the results in this section because these techniques did not demonstrate the best result for any of the molecules tested. The simulation results of all template-based modeling techniques, including MC-Sym and VFoldLA, are presented in S2 Table.

**Table 2.**
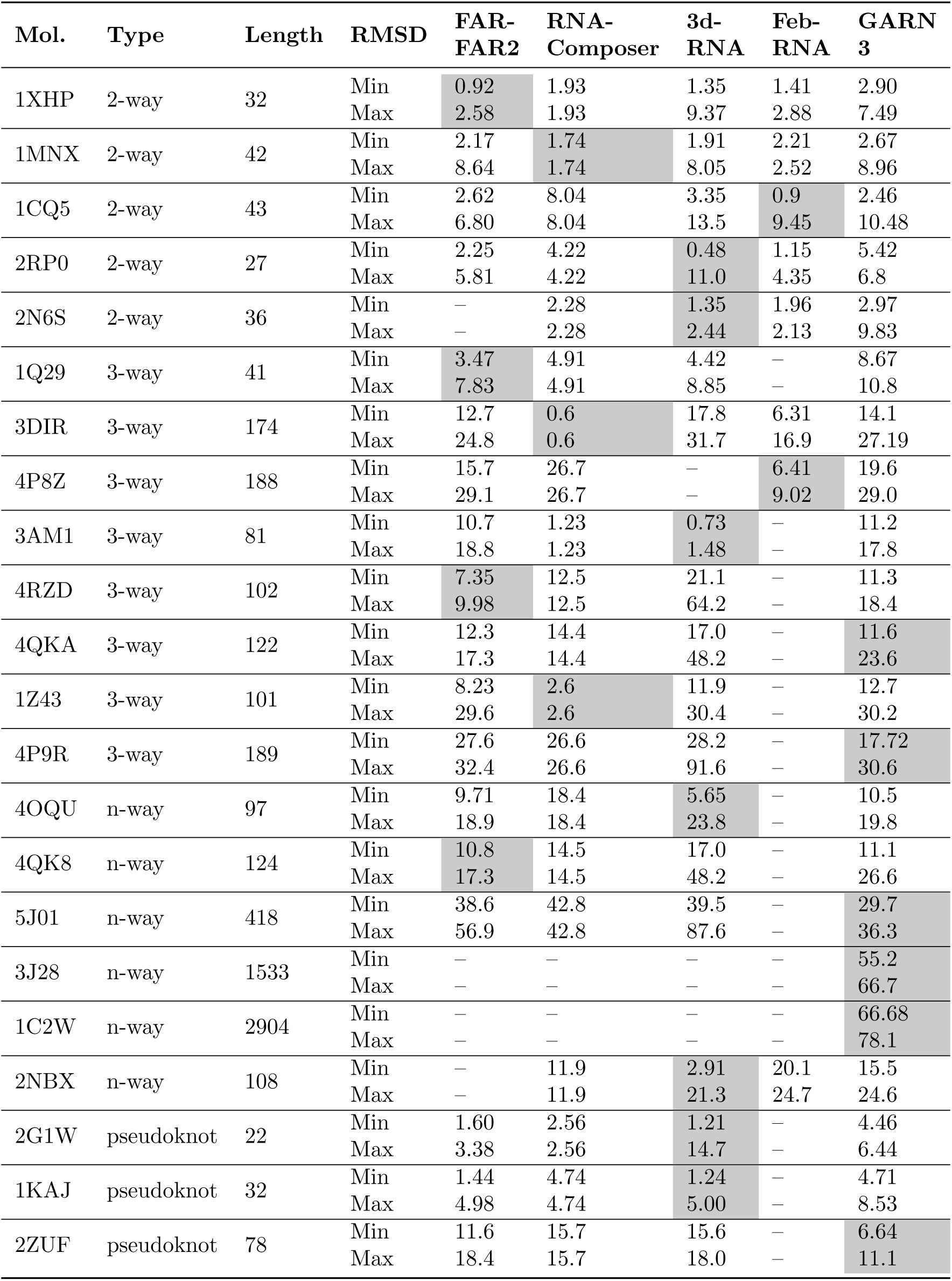
Comparison with template-based modeling techniques. The highlighted values in each row aim to help the visualization of the technique with lower RMSD for the molecule.

| Mol. | Type | Length | RMSD | FAR-FAR2 | RNA-Composer | 3d-RNA | Feb-RNA | GARN 3 |
| --- | --- | --- | --- | --- | --- | --- | --- | --- |
| 1XHP | 2-way | 32 | Min | 0.92 | 1.93 | 1.35 | 1.41 | 2.90 |
|  |  |  | Max | 2.58 | 1.93 | 9.37 | 2.88 | 7.49 |
| 1MNX | 2-way | 42 | Min | 2.17 | 1.74 | 1.91 | 2.21 | 2.67 |
|  |  |  | Max | 8.64 | 1.74 | 8.05 | 2.52 | 8.96 |
| 1CQ5 | 2-way | 43 | Min | 2.62 | 8.04 | 3.35 | 0.9 | 2.46 |
|  |  |  | Max | 6.80 | 8.04 | 13.5 | 9.45 | 10.48 |
| 2RP0 | 2-way | 27 | Min | 2.25 | 4.22 | 0.48 | 1.15 | 5.42 |
|  |  |  | Max | 5.81 | 4.22 | 11.0 | 4.35 | 6.8 |
| 2N6S | 2-way | 36 | Min | – | 2.28 | 1.35 | 1.96 | 2.97 |
|  |  |  | Max | – | 2.28 | 2.44 | 2.13 | 9.83 |
| 1Q29 | 3-way | 41 | Min | 3.47 | 4.91 | 4.42 | – | 8.67 |
|  |  |  | Max | 7.83 | 4.91 | 8.85 | – | 10.8 |
| 3DIR | 3-way | 174 | Min | 12.7 | 0.6 | 17.8 | 6.31 | 14.1 |
|  |  |  | Max | 24.8 | 0.6 | 31.7 | 16.9 | 27.19 |
| 4P8Z | 3-way | 188 | Min | 15.7 | 26.7 | – | 6.41 | 19.6 |
|  |  |  | Max | 29.1 | 26.7 | – | 9.02 | 29.0 |
| 3AM1 | 3-way | 81 | Min | 10.7 | 1.23 | 0.73 | – | 11.2 |
|  |  |  | Max | 18.8 | 1.23 | 1.48 | – | 17.8 |
| 4RZD | 3-way | 102 | Min | 7.35 | 12.5 | 21.1 | – | 11.3 |
|  |  |  | Max | 9.98 | 12.5 | 64.2 | – | 18.4 |
| 4QKA | 3-way | 122 | Min | 12.3 | 14.4 | 17.0 | – | 11.6 |
|  |  |  | Max | 17.3 | 14.4 | 48.2 | – | 23.6 |
| 1Z43 | 3-way | 101 | Min | 8.23 | 2.6 | 11.9 | – | 12.7 |
|  |  |  | Max | 29.6 | 2.6 | 30.4 | – | 30.2 |
| 4P9R | 3-way | 189 | Min | 27.6 | 26.6 | 28.2 | – | 17.72 |
|  |  |  | Max | 32.4 | 26.6 | 91.6 | – | 30.6 |
| 4OQU | n-way | 97 | Min | 9.71 | 18.4 | 5.65 | – | 10.5 |
|  |  |  | Max | 18.9 | 18.4 | 23.8 | – | 19.8 |
| 4QK8 | n-way | 124 | Min | 10.8 | 14.5 | 17.0 | – | 11.1 |
|  |  |  | Max | 17.3 | 14.5 | 48.2 | – | 26.6 |
| 5J01 | n-way | 418 | Min | 38.6 | 42.8 | 39.5 | – | 29.7 |
|  |  |  | Max | 56.9 | 42.8 | 87.6 | – | 36.3 |
| 3J28 | n-way | 1533 | Min | – | – | – | – | 55.2 |
|  |  |  | Max | – | – | – | – | 66.7 |
| 1C2W | n-way | 2904 | Min | – | – | – | – | 66.68 |
|  |  |  | Max | – | – | – | – | 78.1 |
| 2NBX | n-way | 108 | Min | – | 11.9 | 2.91 | 20.1 | 15.5 |
|  |  |  | Max | – | 11.9 | 21.3 | 24.7 | 24.6 |
| 2G1W | pseudoknot | 22 | Min | 1.60 | 2.56 | 1.21 | – | 4.46 |
|  |  |  | Max | 3.38 | 2.56 | 14.7 | – | 6.44 |
| 1KAJ | pseudoknot | 32 | Min | 1.44 | 4.74 | 1.24 | – | 4.71 |
|  |  |  | Max | 4.98 | 4.74 | 5.00 | – | 8.53 |
| 2ZUF | pseudoknot | 78 | Min | 11.6 | 15.7 | 15.6 | – | 6.64 |
|  |  |  | Max | 18.4 | 15.7 | 18.0 | – | 11.1 |

For template-based techniques, considering 2-way molecules, 3dRNA and FebRNA demonstrated good results, performing well in most of the molecules. For 3-way and higher-order junctions (n-way), FARFAR2 performed well and returned good results; in n-way molecules, GARN3 also showed good results, and since the other techniques have some length limitations, 3J28 and 1C2W were only possible to predict by GARN3. When checking the results of deep learning modeling techniques (see S3 Table), trRosettaRNA demonstrated the ability to predict molecules with the best results for most of the molecules (14 out of 22 molecules). Followed by trRosettaRNA, AlphaFold3 also demonstrated good results, followed by GARN3.

When the results from GARN2 and GARN3 are compared, there is a relevant difference. In summary, 18 of the 22 molecules tested demonstrated better results for GARN3. From the molecules better predicted by GARN2, there is no apparent pattern in addition to the structure length, which is small.

### Classification of best molecules

In our simulations, most molecules with 2-way junctions showed better simulation results with the highest maximum distances. In these results, we can take into account the length of the molecules, which are small, having less than 100 nucleotides. The only molecule that had results with the lowest maximum distances was 2RP0, which, despite being among the specific molecules with a 2-way topology, also has a pseudoknot. Likewise, in the molecules with pseudoknots, most of the molecules in the database from this study had the best simulation results with the lowest maximum distances.

In molecules with 3-way junctions, it was possible to verify the opposite, where the molecules were better classified with the lowest maximum distances. This was visible when the molecules are larger and, especially, with larger helices and more nucleotides within these helices. In molecules with n-way junctions, the pattern of smaller distances is also visible. One of the molecules with fewer nucleotides, 2NBX, shows the opposite, possibly due to its smaller size compared to the majority. All results are listed in S3 Fig and S4 Fig.

### Execution time

Another important aspect to check is the execution time, in order to understand if GARN3 maintains the optimized prediction time compared to other techniques in the literature, such as GARN/GARN2 [10, 17]. GARN2 presents a model with a significantly lower quantity of players depending on the molecule; In large molecules, especially the ones with large helices, the 3D model pseudoatoms are more visible than in smaller molecules. Therefore, it is expected that the simulation time of GARN3 increases compared to GARN2, but it is also expected to maintain the efficiency of time compared to other techniques. The difference in the quantity of players between GARN2 and GARN3 models can be seen in S6 Table. After verifying the results, GARN3 demonstrated that it maintains execution time efficiency compared to the other techniques. In fact, the molecules with time-consuming simulations are the longer ones. Other techniques with low execution time are AlphaFold3, RNAComposer, FebRNA and GARN2.

## Discussion

The main objective of this work is to improve the GARN2 technique by adding more players in the helices, enhancing the visualization of the predicted molecules, in our proposed technique named GARN3. To achieve this improvement, we updated the game strategies and introduced a different way of computing the scores with the help of a machine learning regression model. After these improvements, we performed simulations for GARN3 and other techniques in the literature using our set of tests. Compared to GARN2, our technique, GARN3, demonstrated good results where GARN3 predicted the majority of molecules (approximately 80% of the test set) better.

One point of attention after updating the game configuration, in addition to the reduction of granularity in the 3D model, was also the execution time, since it is an important advantage of GARN/GARN2 when compared with other techniques. In our simulations, GARN3 showed an increase in simulation time when the molecules are larger, usually larger than 100 nucleotides, which was expected considering the increase in players. However, the technique showed to be still similar and, in many moments, better when compared to other techniques’ simulation times.

When comparing GARN3 with the other techniques in addition to GARN2, it demonstrated better or equivalent results in our tests. In Fig 10, the simulation results for the test set in this paper are presented for all techniques that were compared in this work. For these results, some similarity can be seen in terms of the RMSD calculation when grouping the results of each of the molecules. Compared with template-based techniques, the differences were more visible in molecules with 3-way junctions and also in n-way junctions with less than 200 nucleotides. Compared to deep learning modeling techniques, GARN3 shows similar results to AlphaFold3 in our test set, although trRosettaRNA demonstrated better predictions for almost all molecules.

**Fig 10.**
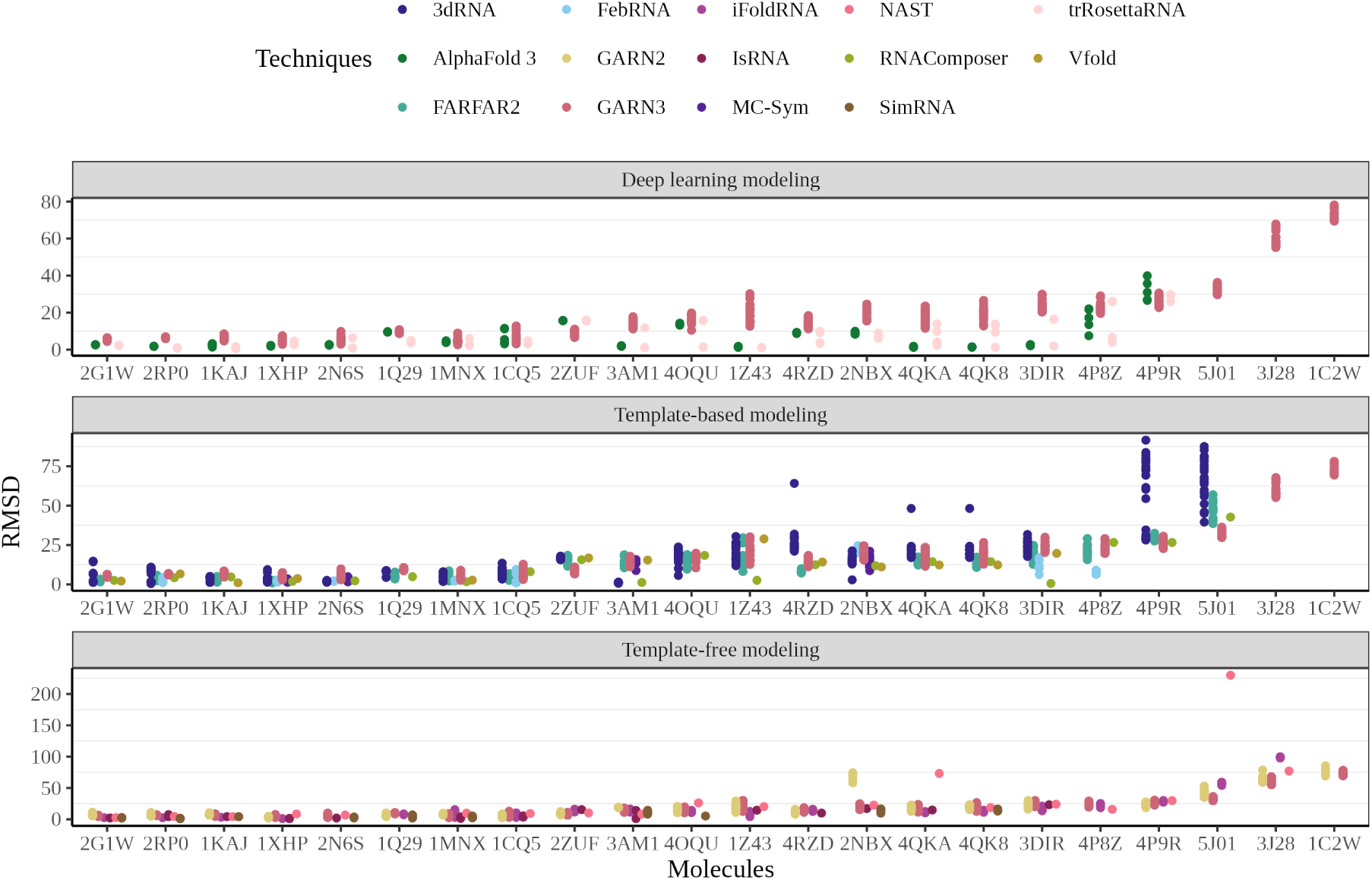
Comparison with other techniques, considering the entire test set, where the results are split by the technique’s simulation type (template-based, template-free, and deep learning). GARN3 is present in the 3 categories, only to compare with the other techniques. The result from GARN3 in this graph represents the best of the sampling, considering both EXP3 and UCB algorithms.

Considering the scoring function, the dynamic distances obtained by the machine learning prediction model enhance the usability of the technique once it reduces the necessity of manually setting the properties that depend on the SSE type. Also, as mentioned before, it predicts some molecules that cannot be predicted by the scoring function with static parameters.

In the methodology section, the usage of maximum distance in the structure is mentioned as the sorting strategy when choosing the best predictions. This method, which is used and validated in GARN2, brought good results to the GARN3 simulations. In some of the molecules, the best results are close to the highest maximum distance, whereas others are close to the lowest maximum. In our simulations, most of the best simulations with the lowest maximum distances are seen in molecules with pseudoknots and also in molecules with 3-way or higher-order junctions. Another pattern found was the best simulations with the highest maximum distances, mostly seen in smaller molecules with 2-way junctions. There is an example in Fig. 11, where the 3DIR molecule has one of the best-evaluated structures with the highest maximum distance, and 3AM1 has one of the best samples with the lowest maximum distance.

**Fig 11.**
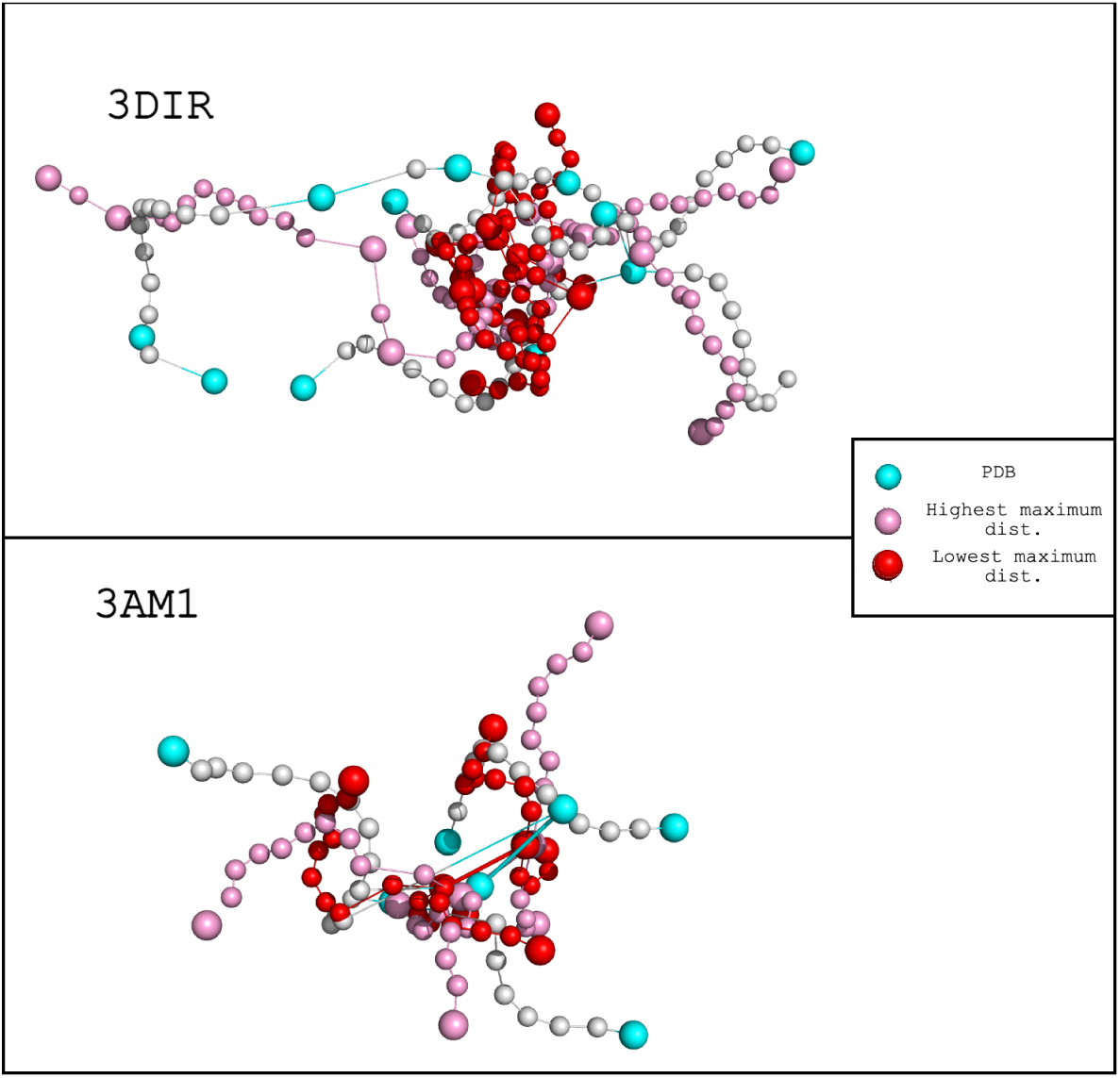
Sorting strategy used in the molecules. The native structure is compared to both the highest and lowest maximum distances. In our simulations, 3DIR has one of the best-evaluated structures with the highest maximum distance, whereas 3AM1 is the opposite, with one of the best samplings from the lowest maximum distance.

A limitation that was not explored here is the combination of global and local scores, since GARN2 and GARN3 compute the scores locally at each node. This is the reason why we decided to use the previously validated sorting criteria in GARN2, which also showed similar results in this study. Another limitation that remains is the quantity of pseudoatoms, which in the GARN model are the players. The technique presents a coarse-grained model, which can be outdated compared to newer techniques that present near-native 3D models in their predictions. Nevertheless, GARN3, just like GARN and GARN2, runs in local machines, relatively fast (as seen before, when compared to some techniques presented in this paper) and without library dependencies, and there is no need for a dedicated GPU (Graphic Processing Unit).

## Conclusion

The results presented GARN3, a coarse-grained approach technique, with good results compared to other previous and also recent techniques. We improved the scoring function used in GARN2 by using a machine learning regression model to retrieve the parameters, which makes the scoring more reliable since we don’t need to manually calculate the parameters, as is done in the previous versions of GARN [10, 17]. The final 3D structure presents more pseudoatoms, having a better approximation to native structures. The simulations using the test set in this study showed interesting results, where GARN3 demonstrated results similar to other template-free modeling techniques. In addition, GARN3 also maintains the efficiency in large-molecule simulations, similar to GARN2, considering that this is still a limitation for many recent techniques. During our tests, we can see that the GARN3 simulations for large molecules showed more approximation to native structures for most of the molecules in the set. During the simulations, it was also visible that a short execution time was performed for most of the molecules.

The GARN3 technique is available at https://github.com/jhonatans01/garn3, written and executable in Java. The repository contains the GARN3 file executable by any Java Virtual Machine (for Java version 17 or higher), as well as a list of instructions for all available functions and parameters.

## Acknowledgments

We thank Mélanie Boudard for providing the GARN2 source code that served as the basis for this work.

## Supporting information

[**S1 Appendix. Molecules used in the analysis.**] All molecule IDs that were used for analysis, both to create the knowledge-based potential and to generate the machine learning prediction model.: 3IAB, 3P59, 1MT4, 1VBX, 1ATO, 1J1U, 1QRS, 1Z43, 2DU4, 2MQT, 1HS4, 3DIS, 2RPT, 1U0B, 3MUT, 3F2Q, 2B7G, 2LA9, 1EUY, 2LV0, 1K6G, 2ES5, 3OL9, 3B31, 4R4V, 3IYR, 1VBY, 157D, 2MIS, 1A4D, 1QF6, 3GOG, 2MER, 2DLC, 2ZH9, 2F88, 1OW9, 3E5C, 1MNX, 1XWU, 3NPN, 1RNK, 1K5I, 3DIG, 3D2G, 1EBR, 3D0X, 4FEO, 2RD2, 4FEN, 3MUV, 4QLN, 1EVV, 2KDQ, 1B23, 4CQN, 1EBS, 4FRG, 3DIQ, 3SKT, 1ZO1, 2DU6, 3IYQ, 2O3V, 1SJ4, 1FFY, 2ZH8, 1VBZ, 1MWL, 1F27, 3GAO, 2M21, 1DRZ, 2M1O, 1FIR, 4GMA, 1A4T, 1A51, 1FEQ, 1SJF, 2F8K, 2OEU, 3IZY, 1QRU, 1T4L, 1XWP, 1HS2, 3E5F, 2HOP, 1Y27, 2LBS, 3F2W, 3MUR, 4FEJ, 2B63, 3CW5, 2LBR, 1Y26, 3DD2, 2Y8Y, 2KRY, 3MOJ, 2ESE, 1L2X, 1L1C, 4R4P, 429D, 1HS3, 3L0U, 1QRT, 2DU3, 2DVI, 2O32, 1QFQ, 1TXS, 4WFL, 28SR, 3GOT, 1SJ3, 2AKE, 1HS1, 2OIU, 3E5E, 1E95, 2G1G, 1U3K, 3D2V, 3F2T, 2B6G, 3F30, 2LBQ, 3CW6, 2L6I, 1NEM, 3NVI, 1R4H, 3DHS, 1L1W, 1EOR, 2RP1, 2JWV, 2OE6, 1VTQ, 3LQX, 4P95, 2FY1, 3GCA, 1DUL, 3BWP, 1Q8N, 2M5U, 1SY4, 1SZY, 3BNR, 2A9L, 357D, 2QH2, 1SCL, 2ZUE, 2XNZ, 2O81, 3IRW, 2GOZ, 3CJZ, 3MJB, 3DW7, 2R93, 3TWH, 1BZ3, 480D, 3OK2, 1IK1, 1LU3, 1BZ2, 2L3C, 3DW6, 3ZD3, 2PN3, 3EGZ, 1ZIG, 2A64, 2QH3, 2ANN, 2DR2, 2O43, 3BNS, 1QTQ, 2HGH, 1AKX, 4XW7, 1O0B, 1F79, 3GER, 4WCP, 2MXL, 422D, 1J4Y, 3B4B, 4L81, 3T4B, 2EVY, 2N2P, 3MJA, 2K4C, 1BYJ, 1YMO, 1LVJ, 2KOC, 2PXK, 1KPZ, 3F4G, 1ESY, 3DW5, 1NTA, 2G5K, 1WTT, 3IQN, 1XSH, 2O83, 1JOX, 1MFQ, 1J7T, 1Q75, 3GES, 1TRA, 1O0C, 5DI2, 5DH6, 1CSL, 1O15, 1AFX, 2QH4, 3LA5, 2DR5, 2O44, 1QU3, 2ZY6, 3BNT, 1NYB, 2YDH, 2RO2, 3A3A, 1BZT, 1BZU, 2K96, 1LUU, 3FO4, 462D, 2KTZ, 3MJ3, 2KX8, 1T28, 2ZZM, 2DRB, 1QU2, 2O45, 1F6X, 5A17, 2AU4, 2FQN, 1F6Z, 2FK6, 3K1V, 2MXJ, 4TZX, 1WTS, 2LPA, 1RFR, 1NBK, 2R8S, 1UUU, 3FO6, 2U2A, 2PXL, 1N8X, 2NQP, 2LI4, 2K95, 1ESH, 439D, 2KXM, 2KXZ, 1Z30, 2DRA, 2DR7, 5DI4, 4YB0, 1H4Q, 1HOQ, 4KQY, 4NYA, 1QWB, 4B5R, 1OQ0, 1XST, 2JSG, 3IQR, 3FU4, 1KD5, 2G9C, 1KH6, 2K5Z, 2PXV, 4FAX, 2K66, 2L3J, 3DVZ, 1LNG, 2KU0, 1LC6, 1KD4, 2KYE, 1RHT, 1XSU, 1TFN, 2JR4, 1TJZ, 3GS5, 3IGI, 4FXD, 2DR9, 1QWA, 4NYB, 2ANR, 1ZJW, 2KX5, 1K2G, 1U9S, 2GCS, 1LUX, 2PXT, 2PXU, 3Q51, 4A4S, 1KPD, 2LK3, 2PXB, 3SUX, 2LP9, 2N2O, 1EHZ, 2KY0, 3IQP, 2JSE, 4NYC, 2X7N, 2DR8, 1MSY, 1F9L, 2FRL, 1Q9A, 1H4S, 4PHY, 1JZC, 1A3M, 2DET, 1SYZ, 1MFK, 1ME1, 1HWQ, 4R0D, 2XLJ, 1KD3, 1NZ1, 2EUY, 2YIF, 3CGR, 1PJY, 2PXQ, 2PXF, 2PXP, 1KP7, 1U8D, 2A43, 2AB4, 1ZIH, 2H0S, 1ME0, 1F7U, 5C7U, 2M8K, 4LX6, 5DH8, 1AJF, 2M58, 5C7W, 2CD1, 4NYD, 2HEM, 1SLO, 1Z2J, 2JRQ, 2XLI, 1EIY, 4MEH, 2YIE, 2KVN, 2LDZ, 2RLU, 3WFS, 2BE0, 2PXE, 2PXD, 3F4H, 4FB0, 2L2K, 1L9A, 1NUJ, 1KFO, 1XSG, 2JRG, 2H2X, 1ZSE, 397D, 1F7V, 1XJR, 2AP5, 2ZH3, 2HW8, 1TN1, 2QBZ, 3DIM, 2GIP, 3DIZ, 3SD1, 2KRW, 2Y8W, 2LUB, 3OWW, 1EUQ, 4JAH, 1YG3, 3F2X, 4TZY, 3F2Y, 2LC8, 1YSV, 1L3D, 2RPK, 2PJP, 3DIL, 2RRC, 3SKI, 430D, 1FQZ, 1TOB, 4E8Q, 1VC5, 1CX0, 3P4C, 2ZH2, 3NDB, 1FJE, 4Y1M, 1VC7, 1TN2, 4EN5, 2OE8, 3IZ4, 1SER, 1JID, 1S2F, 2HOK, 3GX3, 1D0T, 1S34, 1VY9, 1ROQ, 3DIY, 2GIS, 4ZNP, 3AL0, 2KF0, 3AMU, 2KD8, 4XNR, 1I9V, 2EES, 3AMT, 3SD3, 1U2A, 3DIX, 2ESI, 4FRN, 3DIO, 3NPQ, 1VY8, 1D0U, 3GX2, 2HOJ, 2XEB, 2JXV, 3G4M, 1A60, 2ZH1, 1VC6, 1AM0, 1VOP, 2ZH5, 2XDB, 1ATW, 4E8V, 3R4F, 3GX6, 3NPB, 2OIH, 1K4A, 3S1R, 1I4C, 1YFG, 3Q51, 4FE5, 2EEW, 2B57, 387D, 2EEV, 3MUM, 1I4B, 1YG4, 3D0U, 2Y95, 5BTP, 3DJ0, 3GX7, 2HOO, 2OJ3, 1MNB, 1ATV, 2TPK, 2ZH4, 2ZHB, 2MFD, 1A9L, 2AP0, 1FHK, 1F1T, 2ZH6, 1ZBN, 2O3X, 2TRA, 1ZL3, 2HOM, 3GX5, 1HS8, 1K4B, 1GTR, 3DJ2, 2LUP, 3OVA, 3MXH, 1KXK, 1I46, 466D, 2L5Z, 2EET, 2NUE, 1EXD, 2RE8, 1PBR, 1P5P, 2EEU, 2LA5, 364D, 2KEZ, 2LAC, 1ETF, 3Q50, 4C4Q, 1K8W, 1GTS, 2F4T, 2HOL, 1HQ1, 2A2E, 4E8T, 2ZH7, 1VC0, 2ZHA, 2HVY, 2MF1.

**S1 Fig.**
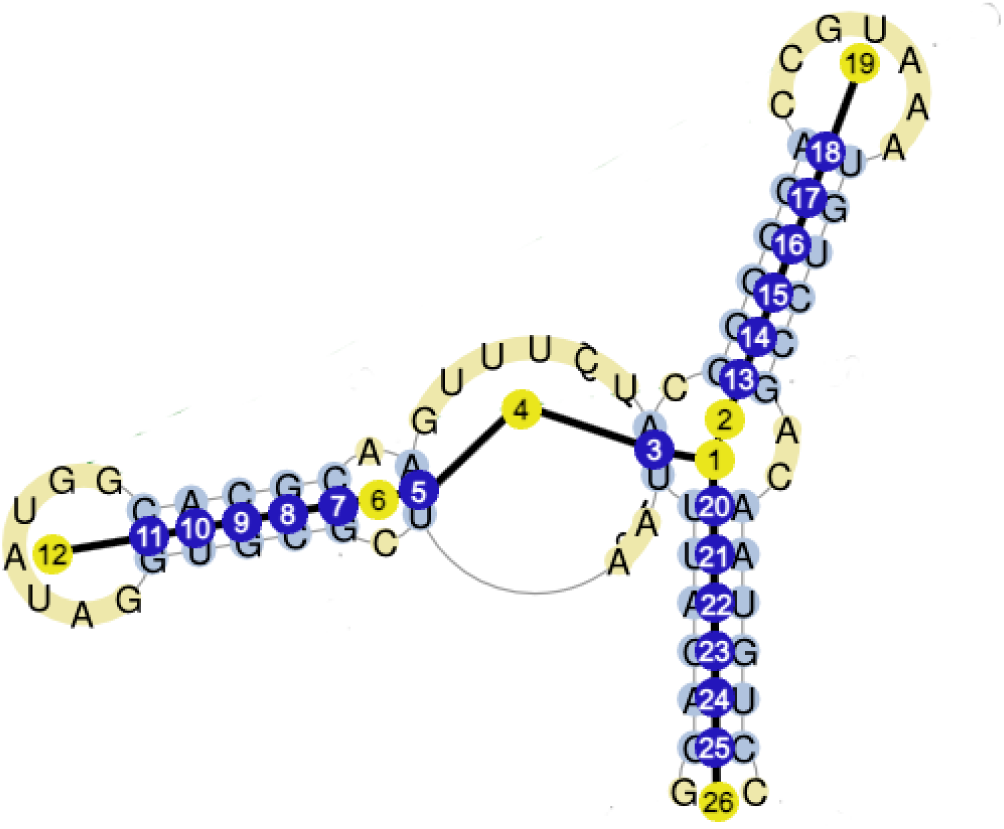
GARN players ordered. First the players of the highest-order junction (in the example, it is a 3-way junction) play, and the order of the other players is defined by applying a depth-first search algorithm, starting the search from the first highest-order junction.

**S2 Fig.**
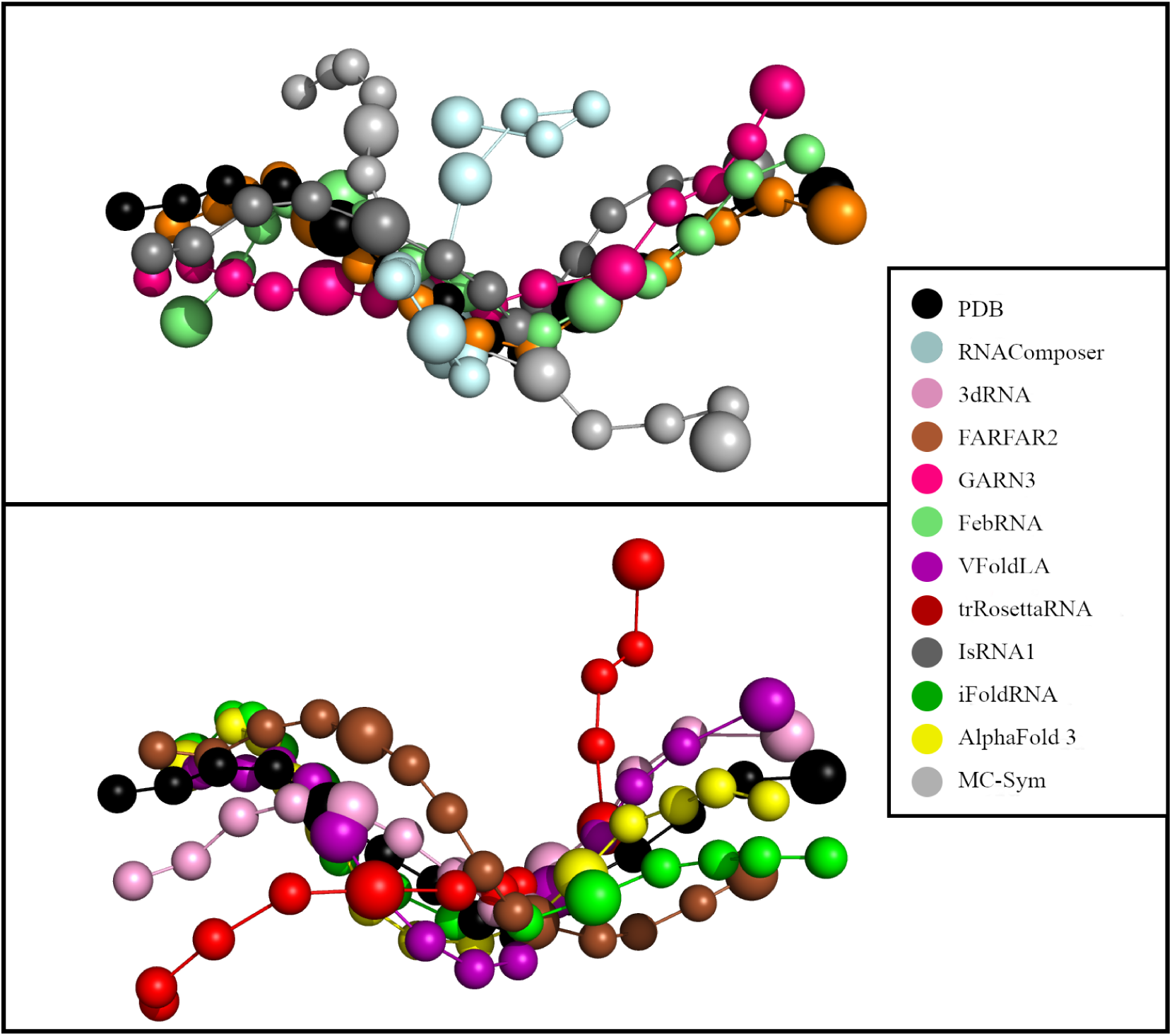
Best simulations for the molecule 1XHP. This figure presents the best simulation for the molecule 1XHP, by each of the techniques tested here. SimRNA is not present because it was not possible to generate the structure. GARN2 is also not present, since it uses a different model with fewer pseudoatoms.

**S1 Table.**
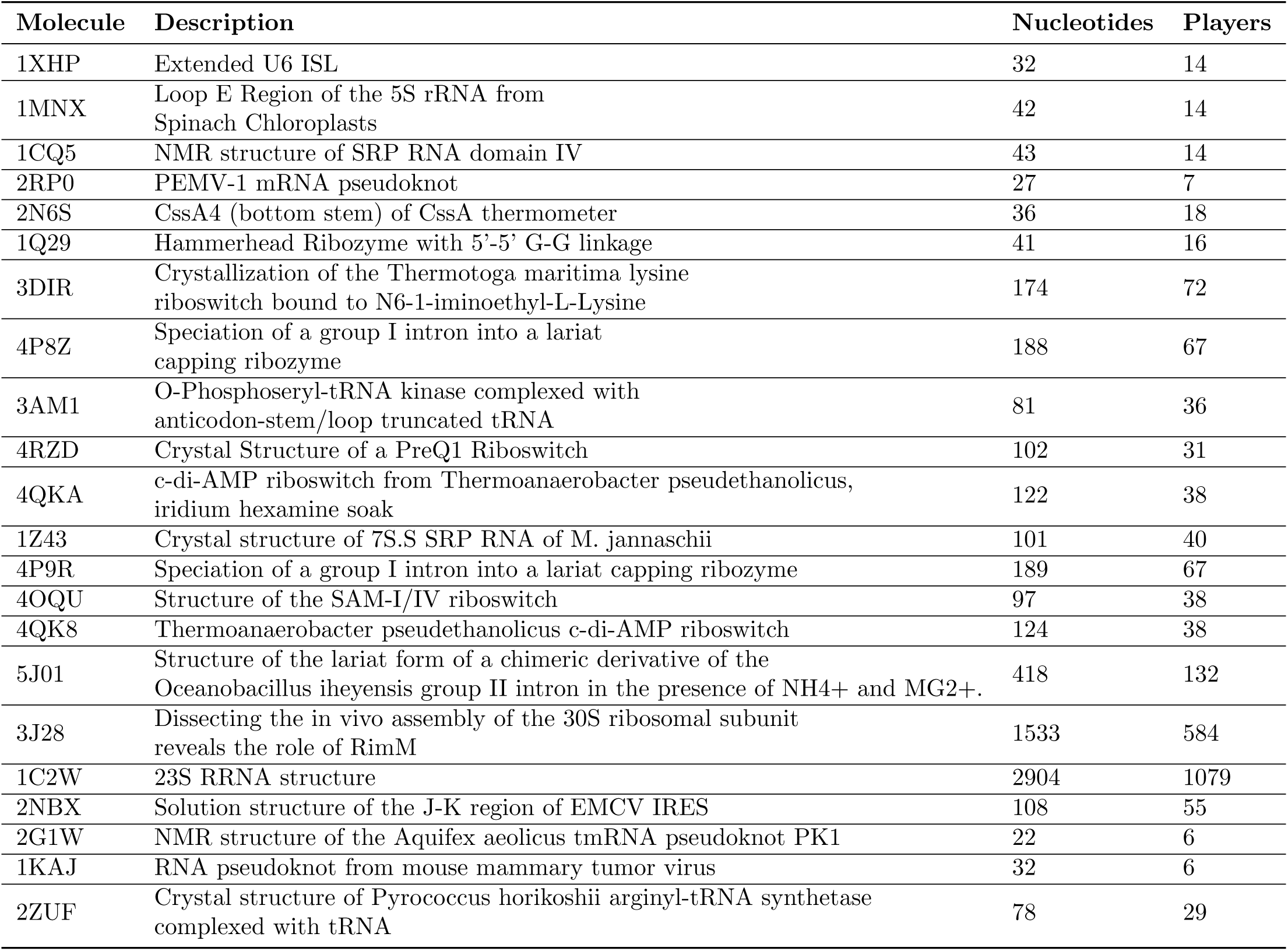
Test set. Molecules used to run the simulations. This test set contains 22 molecules.

**S2 Table.** Template-based techniques simulations results. Comparison of GARN3 with other techniques, considering only template-based techniques.

| Mol. | Type | Length | RMSD | FAR-FAR2 | RNA-Composer | VFold-LA | 3d-RNA | Feb-RNA | MC-Sym | GARN3 |
| --- | --- | --- | --- | --- | --- | --- | --- | --- | --- | --- |
| 1XHP | 2-way | 32 | Min | 0.92 | 1.93 | 3.70 | 1.35 | 1.41 | 1.38 | 2.90 |
|  |  |  | Max | 2.58 | 1.93 | 3.70 | 9.37 | 2.88 | 3.52 | 7.49 |
| 1MNX | 2-way | 42 | Min | 2.17 | 1.74 | 2.72 | 1.91 | 2.21 | – | 2.67 |
|  |  |  | Max | 8.64 | 1.74 | 2.72 | 8.05 | 2.52 | – | 8.96 |
| 1CQ5 | 2-way | 43 | Min | 2.62 | 8.04 | 1.01 | 3.35 | 0.9 | – | 2.46 |
|  |  |  | Max | 6.80 | 8.04 | 1.01 | 13.5 | 9.45 | – | 10.48 |
| 2RP0 | 2-way | 27 | Min | 2.25 | 4.22 | 6.68 | 0.48 | 1.15 | – | 5.42 |
|  |  |  | Max | 5.81 | 4.22 | 6.68 | 11.0 | 4.35 | – | 6.8 |
| 2N6S | 2-way | 36 | Min | – | 2.28 | – | 1.35 | 1.96 | 1.62 | 2.97 |
|  |  |  | Max | – | 2.28 | – | 2.44 | 2.13 | 4.86 | 9.83 |
| 1Q29 | 3-way | 41 | Min | 3.47 | 4.91 | – | 4.42 | – | – | 8.67 |
|  |  |  | Max | 7.83 | 4.91 | – | 8.85 | – | – | 10.8 |
| 3DIR | 3-way | 174 | Min | 12.7 | 0.6 | 19.7 | 17.8 | 6.31 | – | 14.1 |
|  |  |  | Max | 24.8 | 0.6 | 19.7 | 31.7 | 16.9 | – | 27.19 |
| 4P8Z | 3-way | 188 | Min | 15.7 | 26.7 | – | – | 6.41 | – | 19.6 |
|  |  |  | Max | 29.1 | 26.7 | – | – | 9.02 | – | 29.0 |
| 3AM1 | 3-way | 81 | Min | 10.7 | 1.23 | 15.5 | 0.73 | – | – | 11.2 |
|  |  |  | Max | 18.8 | 1.23 | 15.5 | 1.48 | – | – | 17.8 |
| 4RZD | 3-way | 102 | Min | 7.35 | 12.5 | 14.2 | 21.1 | – | – | 11.3 |
|  |  |  | Max | 9.98 | 12.5 | 14.2 | 64.2 | – | – | 18.4 |
| 4QKA | 3-way | 122 | Min | 12.3 | 14.4 | 12.3 | 17.0 | – | – | 11.6 |
|  |  |  | Max | 17.3 | 14.4 | 12.3 | 48.2 | – | – | 23.6 |
| 1Z43 | 3-way | 101 | Min | 8.23 | 2.6 | 28.9 | 11.9 | – | – | 12.7 |
|  |  |  | Max | 29.6 | 2.6 | 28.9 | 30.4 | – | – | 30.2 |
| 4P9R | 3-way | 189 | Min | 27.6 | 26.6 | – | 28.2 | – | – | 17.72 |
|  |  |  | Max | 32.4 | 26.6 | – | 91.6 | – | – | 30.6 |
| 4OQU | n-way | 97 | Min | 9.71 | 18.4 | – | 5.65 | – | – | 10.5 |
|  |  |  | Max | 18.9 | 18.4 | – | 23.8 | – | – | 19.8 |
| 4QK8 | n-way | 124 | Min | 10.8 | 14.5 | 12.4 | 17.0 | – | – | 11.1 |
|  |  |  | Max | 17.3 | 14.5 | 12.4 | 48.2 | – | – | 26.6 |
| 5J01 | n-way | 418 | Min | 38.6 | 42.8 | – | 39.5 | – | – | 29.7 |
|  |  |  | Max | 56.9 | 42.8 | – | 87.6 | – | – | 36.3 |
| 3J28 | n-way | 1533 | Min | – | – | – | – | – | – | 55.2 |
|  |  |  | Max | – | – | – | – | – | – | 66.7 |
| 1C2W | n-way | 2904 | Min | – | – | – | – | – | – | 66.68 |
|  |  |  | Max | – | – | – | – | – | – | 78.1 |
| 2NBX | n-way | 108 | Min | – | 11.9 | 11.1 | 2.91 | 20.1 | 8.75 | 15.5 |
|  |  |  | Max | – | 11.9 | 11.1 | 21.3 | 24.7 | 21.1 | 24.6 |
| 2G1W | pseudo-knot | 22 | Min | 1.60 | 2.56 | 2.15 | 1.21 | – | – | 4.46 |
|  |  |  | Max | 3.38 | 2.56 | 2.15 | 14.7 | – | – | 6.44 |
| 1KAJ | pseudo-knot | 32 | Min | 1.44 | 4.74 | 1.01 | 1.24 | – | – | 4.71 |
|  |  |  | Max | 4.98 | 4.74 | 1.01 | 5.00 | – | – | 8.53 |
| 2ZUF | pseudo-knot | 78 | Min | 11.6 | 15.7 | 16.8 | 15.6 | – | – | 6.64 |
|  |  |  | Max | 18.4 | 15.7 | 16.8 | 18.0 | – | – | 11.1 |

**S3 Table.** Deep learning based techniques simulations results. Comparison of GARN3 with other techniques, considering only techniques based on deep learning approach.

| Mol. | Type | Length | RMSD | AlphaFold3 | trRosettaRNA | GARN3 |
| --- | --- | --- | --- | --- | --- | --- |
| 1XHP | 2-way | 32 | Min | 1.82 | 2.75 | 2.90 |
|  |  |  | Max | 2.36 | 4.72 | 7.49 |
| 1MNX | 2-way | 42 | Min | 3.79 | 2.25 | 2.67 |
|  |  |  | Max | 4.81 | 6.09 | 8.96 |
| 1CQ5 | 2-way | 43 | Min | 3.12 | 3.14 | 2.46 |
|  |  |  | Max | 11.6 | 4.94 | 10.48 |
| 2RP0 | 2-way | 27 | Min | 1.76 | 0.79 | 5.42 |
|  |  |  | Max | 1.87 | 1.13 | 6.8 |
| 2N6S | 2-way | 36 | Min | 2.37 | 0.86 | 2.97 |
|  |  |  | Max | 2.80 | 6.43 | 9.83 |
| 1Q29 | 3-way | 41 | Min | 9.49 | 3.66 | 8.67 |
|  |  |  | Max | 9.77 | 5.01 | 10.8 |
| 3DIR | 3-way | 174 | Min | 2.07 | 1.99 | 14.1 |
|  |  |  | Max | 2.77 | 16.5 | 27.19 |
| 4P8Z | 3-way | 188 | Min | 7.50 | 3.85 | 19.6 |
|  |  |  | Max | 22.1 | 26.1 | 29.0 |
| 3AM1 | 3-way | 81 | Min | 1.74 | 1.11 | 11.2 |
|  |  |  | Max | 2.19 | 11.8 | 17.8 |
| 4RZD | 3-way | 102 | Min | 8.66 | 3.40 | 11.3 |
|  |  |  | Max | 9.37 | 9.89 | 18.4 |
| 4QKA | 3-way | 122 | Min | 1.26 | 2.48 | 11.6 |
|  |  |  | Max | 1.85 | 14.0 | 23.6 |
| 1Z43 | 3-way | 101 | Min | 1.30 | 1.03 | 12.7 |
|  |  |  | Max | 1.82 | 1.3 | 30.2 |
| 4P9R | 3-way | 189 | Min | 26.6 | 25.9 | 17.72 |
|  |  |  | Max | 39.9 | 29.8 | 30.6 |
| 4OQU | n-way | 97 | Min | 13.2 | 1.47 | 10.5 |
|  |  |  | Max | 14.3 | 15.8 | 19.8 |
| 4QK8 | n-way | 124 | Min | 1.30 | 1.25 | 11.1 |
|  |  |  | Max | 1.56 | 13.8 | 26.6 |
| 5J01 | n-way | 418 | Min | – | – | 29.7 |
|  |  |  | Max | – | – | 36.3 |
| 3J28 | n-way | 1533 | Min | – | – | 55.2 |
|  |  |  | Max | – | – | 66.7 |
| 1C2W | n-way | 2904 | Min | – | – | 66.68 |
|  |  |  | Max | – | – | 78.1 |
| 2NBX | n-way | 108 | Min | 8.36 | 6.18 | 15.5 |
|  |  |  | Max | 9.94 | 9.0 | 24.6 |
| 2G1W | pseudoknot | 22 | Min | 2.57 | 2.21 | 4.46 |
|  |  |  | Max | 2.74 | 2.37 | 6.44 |
| 1KAJ | pseudoknot | 32 | Min | 1.38 | 0.6 | 4.71 |
|  |  |  | Max | 3.27 | 1.59 | 8.53 |
| 2ZUF | pseudoknot | 78 | Min | 15.7 | 15.6 | 6.64 |
|  |  |  | Max | 15.8 | 15.9 | 11.1 |

**S4 Table.** Simulations using both regret minimization algorithms. Comparison of GARN3 simulations using UCB and EXP3 algorithms.

| Molecule | Type | Length | Players | RMSD | EXP3 | UCB |
| --- | --- | --- | --- | --- | --- | --- |
| 1XHP | 2-way | 32 | 14 | Min | 2.41 | 1.85 |
|  |  |  |  | Max | 9.2 | 8.26 |
| 1MNX | 2-way | 42 | 14 | Min | 3.48 | 2.67 |
|  |  |  |  | Max | 9.53 | 8.96 |
| 1CQ5 | 2-way | 43 | 14 | Min | 3.9 | 2.46 |
|  |  |  |  | Max | 10.76 | 10.48 |
| 2RP0 | 2-way | 27 | 7 | Min | 5.42 | 5.6 |
|  |  |  |  | Max | 6.8 | 6.8 |
| 2N6S | 2-way | 36 | 18 | Min | 2.97 | 3.0 |
|  |  |  |  | Max | 9.83 | 9.77 |
| 1Q29 | 3-way | 41 | 16 | Min | 8.67 | 9.34 |
|  |  |  |  | Max | 10.8 | 9.63 |
| 3DIR | 3-way | 174 | 72 | Min | 14.1 | 16.57 |
|  |  |  |  | Max | 27.19 | 29.96 |
| 4P8Z | 3-way | 188 | 67 | Min | 19.6 | 20.98 |
|  |  |  |  | Max | 29.0 | 28.67 |
| 3AM1 | 3-way | 81 | 36 | Min | 11.2 | 10.30 |
|  |  |  |  | Max | 17.8 | 18.49 |
| 4RZD | 3-way | 102 | 31 | Min | 11.3 | 11.3 |
|  |  |  |  | Max | 18.4 | 19.34 |
| 4QKA | 3-way | 122 | 38 | Min | 11.6 | 13.24 |
|  |  |  |  | Max | 23.6 | 25.97 |
| 1Z43 | 3-way | 101 | 40 | Min | 12.7 | 13.07 |
|  |  |  |  | Max | 30.2 | 29.36 |
| 4P9R | 3-way | 189 | 67 | Min | 17.72 | 19.77 |
|  |  |  |  | Max | 30.6 | 32.20 |
| 4OQU | n-way | 97 | 38 | Min | 10.5 | 11.5 |
|  |  |  |  | Max | 19.8 | 20.6 |
| 4QK8 | n-way | 124 | 38 | Min | 11.1 | 8.0 |
|  |  |  |  | Max | 26.6 | 20.62 |
| 5J01 | n-way | 418 | 132 | Min | 29.7 | – |
|  |  |  |  | Max | 36.3 | – |
| 3J28 | n-way | 1533 | 527 | Min | 55.2 | – |
|  |  |  |  | Max | 66.7 | – |
| 1C2W | n-way | 2904 | 1079 | Min | 66.68 | – |
|  |  |  |  | Max | 78.1 | – |
| 2NBX | n-way | 108 | 55 | Min | 15.5 | 16.57 |
|  |  |  |  | Max | 24.6 | 28.78 |
| 2G1W | pseudoknot | 22 | 6 | Min | 4.46 | 4.35 |
|  |  |  |  | Max | 6.44 | 6.36 |
| 1KAJ | pseudoknot | 32 | 6 | Min | 4.71 | 4.13 |
|  |  |  |  | Max | 8.53 | 10.13 |
| 2ZUF | pseudoknot | 78 | 29 | Min | 6.64 | 6.14 |
|  |  |  |  | Max | 11.1 | 10.42 |

**S3 Fig.**
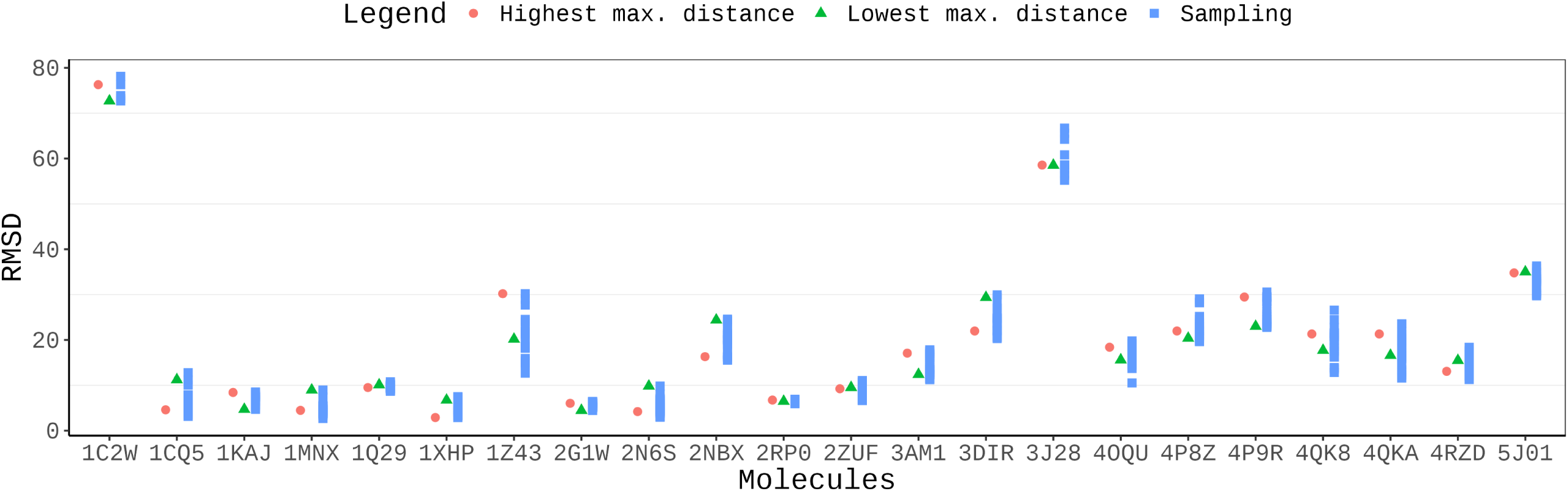
Maximum distance of GARN3 sampling for the test set. This plot presents the highest maximum and lowest maximum distances between the nodes, for each of the molecules in the test set.

**S4 Fig.**
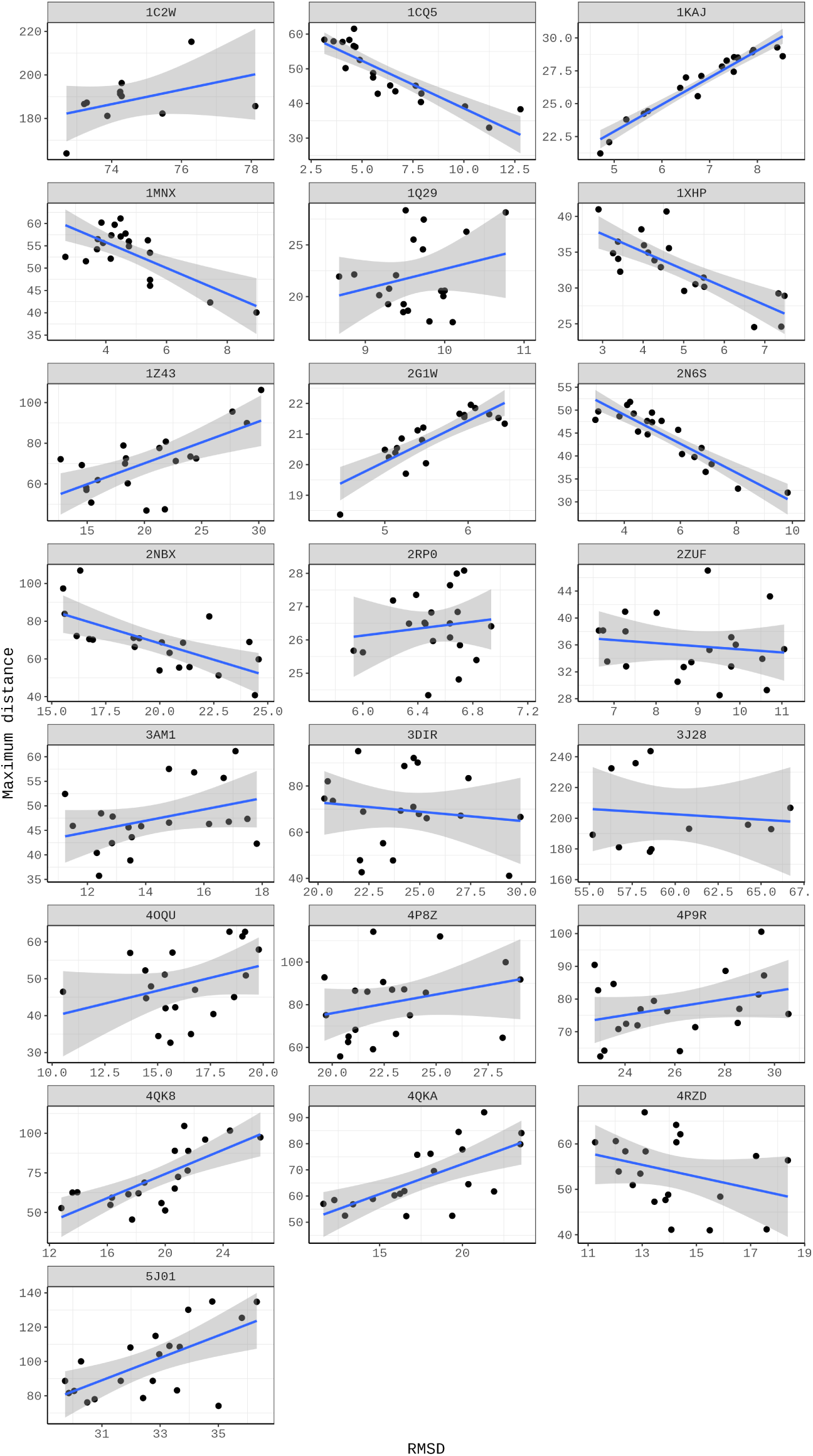
Relation between RMSD and the maximum distance of GARN3 sampling, for the given test set. This plot presents the RMSD and maximum distances between the nodes for each of the molecules in the test set.

**S5 Table.** Simulation time of the molecules from the test set. The asterisk sign (*) by the side of the technique name represents that the simulations were run on a local machine, and the others were on its dedicated web server. The VFoldLA is not present because, even when providing different email addresses from different domains, the web server did not send any notification when the simulations finished.

| Molecule | # | FAR-FAR2 | iFold-RNA | NAST* | RNA-Compo-ser | SimRNA | 3dRNA | isRNA1 | Feb-RNA* | Alpha-Fold | MC-Sym | trRo-setta-RNA | GARN 2* | GARN 3* |
| --- | --- | --- | --- | --- | --- | --- | --- | --- | --- | --- | --- | --- | --- | --- |
| 1XHP | 32 | ~16m | ~13m | ~1m | ~1m | NA | ~47m | ~2h | ~1m | ~3m | ~25m | ~10m | ~1m | ~1m |
| 1MNX | 42 | ~19m | ~18m | ~1m | ~1m | ~1h25 | ~52m | ~4h | ~1m | ~5m | NA | ~10m | ~1m | ~1m |
| 1CQ5 | 43 | ~20m | ~27m | ~1m | ~1m | NA | ~52m | ~4h | ~1m | ~5m | NA | ~1m | ~1m | ~1m |
| 2RP0 | 27 | ~22m | ~13m | ~1m | ~1m | ~1h | ~1h22 | ~2h | ~1m | ~3m | NA | ~1m | ~1m | ~1m |
| 2N6S | 36 | NA | NA | ~1m | ~1m | ~1h20 | ~44m | ~1h | ~1m | ~3m | ~30m | ~1m | NA | ~1m |
| 1Q29 | 41 | ~34m | ~19m | ~1m | ~1m | ~10m | ~2h | NA | ~1m | ~5m | NA | ~10m | ~1m | ~1m |
| 3DIR | 174 | ~3h13 | ~2h08 | ~1m | ~1m | ~12h26 | ~10h12 | ~1d | ~4m | ~10m | NA | ~8m | ~1m | ~14m |
| 4P8Z | 188 | ~6d10h | ~2h13 | ~1m | ~9m | ~13h05 | ~2h18 | NA | ~2m | ~11m | NA | ~1h | NA | ~12m |
| 3AM1 | 81 | ~35m | ~1h30 | ~1m | ~3m | ~1d7h | ~1h07 | ~6h30 | ~1m | ~9m | ~32m | ~46m | ~1m | ~15m |
| 4RZD | 102 | ~46m | ~48m | ~1m | ~3m | ~4h49 | ~6h45 | ~13h20 | ~1m | ~8m | NA | ~26m | ~1m | ~1m |
| 4QKA | 122 | ~1h58 | ~1h52 | ~1m | ~5m | ~6h | ~6h20 | ~12h | ~1m | ~6m | NA | ~54m | ~1m | ~2m |
| 1Z43 | 101 | ~52m | ~1h | ~1m | ~4m | NA | ~1h42 | ~8h16 | ~1m | ~10m | NA | ~42m | ~1m | ~3m |
| 4P9R | 189 | ~6h59 | ~2h15 | ~1m | ~9m | NA | ~11h22 | NA | ~2m | ~10m | NA | ~1h24 | ~1m | ~13m |
| 4OQU | 97 | ~1h42 | ~50m | ~1m | ~3m | ~3h12 | ~4h39 | NA | ~1m | ~9m | NA | ~54m | ~1m | ~2m |
| 4QK8 | 124 | ~10h29 | ~1h15 | ~1m | ~5m | NA | ~6h41 | NA | ~1m | ~7m | NA | ~58m | ~1m | ~2m |
| 5J01 | 418 | ~12h34 | ~5h33 | ~10m | ~24m | NA | ~1d20h | NA | NA | NA | NA | ~2h46 | ~1m | ~9m |
| 3J28 | 1533 | NA | ~1d9h | ~14h05 | NA | NA | NA | NA | NA | NA | NA | NA | ~1m | ~2h51 |
| 1C2W | 2904 | NA | NA | NA | NA | NA | NA | NA | NA | NA | NA | NA | ~1m | ~3h18 |
| 2NBX | 108 | NA | NA | ~1m | ~4m | ~1d6h | ~2h04 | ~9h44 | ~2m | ~15m | ~33m | ~38m | ~1m | ~9m |
| 2G1W | 22 | ~15m | ~8m | ~1m | ~1m | ~35m | ~1h24 | ~3h13 | ~1m | ~2m | NA | ~21m | ~1m | ~1m |
| 1KAJ | 32 | ~20m | ~12m | ~1m | ~1m | ~32m | ~2h05 | ~3h48 | ~1m | ~3m | NA | ~24m | ~1m | ~1m |
| 2ZUF | 78 | ~7h30 | ~36m | ~1m | ~2m | ~2h40 | ~1h14 | ~7h34 | ~1m | ~7m | NA | ~1h33 | ~1m | ~1m |

**S6 Table.** Players in GARN2 and GARN3 models. Quantity of players used when simulating the molecules in the test set, considering GARN2 and GARN3 models.

| Molecule | Nucleotides | Players<br>GARN2 | Players<br>GARN3 |
| --- | --- | --- | --- |
| 1XHP | 32 | 6 | 14 |
| 1MNX | 42 | 5 | 14 |
| 1CQ5 | 43 | 6 | 14 |
| 2RP0 | 27 | 3 | 7 |
| 2N6S | 36 | 4 | 18 |
| 1Q29 | 41 | 8 | 16 |
| 3DIR | 174 | 32 | 72 |
| 4P8Z | 188 | NA | 67 |
| 3AM1 | 81 | 15 | 36 |
| 4RZD | 102 | 13 | 31 |
| 4QKA | 122 | 20 | 38 |
| 1Z43 | 101 | 18 | 40 |
| 4P9R | 189 | 30 | 67 |
| 4OQU | 97 | 18 | 38 |
| 4QK8 | 124 | 20 | 38 |
| 5J01 | 418 | 66 | 132 |
| 3J28 | 1533 | 310 | 584 |
| 1C2W | 2904 | 505 | 1079 |
| 2NBX | 108 | 24 | 55 |
| 2G1W | 22 | 3 | 6 |
| 1KAJ | 32 | 3 | 6 |
| 2ZUF | 78 | 13 | 29 |

